# Organophosphorus pesticide and nerve agent surrogate metabolism by human CYP3A4

**DOI:** 10.64898/2026.04.23.720309

**Authors:** Pratik Shriwas, Abigail M. Noonchester, Brian T. Scarpitti, Andre Revnew, Thomas R. Lane, Sean Ekins, Christopher M. Hadad, Craig A. McElroy

## Abstract

Of the cytochrome P450 enzymes, CYP3A4 is the most abundant isoform in the human liver, and this enzyme plays a dominant role in the metabolism of a wide range of clinical drugs and xenobiotics. Previous studies have demonstrated that CYP3A4 participates in the oxidative metabolism of several organophosphorus (OP) pesticides involving both thion (P=S) and oxon (P=O) forms. In the present study, we evaluated the capacity of CYP3A4 to metabolize a structurally diverse set of OP compounds using LC-MS/MS methods and assessed their potential to inhibit CYP3A4 activity using previously developed pFlour50 fluorogenic assay. Our results demonstrate that CYP3A4 preferentially metabolizes thions, as compared to oxons, and several OP compounds were also found to inhibit CYP3A4 activity in a time-dependent manner. To gain further mechanistic structural insight into the CYP3A4-OP interactions, molecular docking studies were performed using a crystal structure of CYP3A4 (PDB ID: 3NXU). Linear correlation analysis between *in silico* parameters like molecular weight or binding energy correlated with experimental data including inhibition data for 10 or 30 minutes or the LC-MS/MS data showing the degradation at 1 or 2 hours showed moderate but significant correlation. Soman surrogate PiMP, and cyclosarin surrogate CMP, were both effectively metabolized by CYP3A4, while docking of these surrogates and authentic agents with CYP3A4 receptor revealed very similar binding poses and interactions. Collectively, these findings highlight the important role of CYP3A4 in OP metabolism and support the potential of integrating experimental and *in silico* data to predict CYP3A4-mediated metabolism of existing and emerging OP compounds, including those of toxicological and chemical warfare relevance.

## Introduction

Organophosphorus (OP) compounds are highly toxic chemicals that exert their biological effects primarily through covalent inhibition of acetylcholinesterase (AChE), leading to excessive accumulation of acetylcholine and disruption of cholinergic neurotransmission. As a result, exposure to OP compounds can produce severe acute toxicity as well as long-term neurological and systemic effects ^1^. Intentional ingestion of OP pesticides represents one of the leading causes of suicide globally, accounting for approximately 150,000–200,000 deaths annually, or nearly 15% of all suicide-related fatalities worldwide ^2–4^. In addition to deliberate exposure, the expanding use of OP pesticides in agriculture has significantly increased the risk of accidental and occupational poisoning. Global pesticide applications exceeded 3.7 million metric tons in 2022 (*Global pesticide consumption 1990-2022*, accessed 27 July 2025), and it is estimated that nearly 25 million agricultural workers experience unintentional pesticide poisoning each year ^6^. Chronic, low-dose exposure through pesticide residues on food commodities such as fruits and vegetables ^7–9^ further contributes to population-wide exposure and has been associated with adverse health outcomes including neurotoxicity and cancer ^10^.

Beyond agricultural settings, OP compounds pose a major threat as chemical warfare nerve agents (CWNAs), which are among the most potent neurotoxic agents known. Several high-profile incidents over the past decade have demonstrated the continued use of OP-based nerve agents in targeted and mass exposure events. These include the sarin attacks in Syria in 2013 ^11^, the VX-mediated assassination of Kim Jong Nam in Malaysia in 2017 ^12^, the Novichok nerve agent poisoning in Salisbury, UK, in 2018 that resulted in the death of Dawn Sturgess, and the poisoning of Alexei Navalny in Russia in 2020 ^13^. Collectively, these recent events highlight the urgent need to better understand endogenous mechanisms that contribute to OP detoxification and clearance in humans.

Structurally, OP compounds are classified as either oxons (P=O) or thions (P=S), depending on the atom double-bonded to the phosphorus center ^14^. CWNAs exist exclusively in the oxon form, whereas many OP pesticides are applied as thions that require metabolic bioactivation via oxidative desulfuration to generate the toxic oxon metabolite responsible for AChE inhibition ^15^. Current medical countermeasures for OP poisoning rely on antimuscarinic agents, such as atropine, in combination with AChE reactivators, including pralidoxime, obidoxime or HI-6 ^16,17^. While these treatments are effective in mitigating peripheral toxicity, their limited penetration across the blood–brain barrier reduces their efficacy within the central nervous system (CNS). Although newer reactivators, such as quinone methide precursor compounds, show promise in addressing these limitations and improving CNS protection, they remain under development and have not yet received clinical approval ^18,19^. Consequently, there is a continued need for alternative or complementary strategies to reduce OP toxicity, particularly within the CNS.

Cytochrome P450 (CYP) enzymes constitute a major defense system against xenobiotics by catalyzing phase I oxidative metabolism of drugs, environmental chemicals, and endogenous substrates. Among these enzymes, CYP3A4 is the most abundant CYP isoform in the adult human liver and intestine, and this enzyme plays a dominant role in xenobiotic clearance ^20^. CYP3A4 alone is estimated to contribute to the metabolism of approximately 30–40% of all clinically used drugs, reflecting its unusually large and flexible active site capable of accommodating structurally diverse substrates ^21,22^. In addition to pharmaceuticals, CYP3A4 has been implicated in the metabolism of multiple classes of pesticides and environmental toxicants ^23–26^, suggesting a potential role as an endogenous bioscavenger for OP compounds.

Recent work from our group demonstrated that human liver microsomes (HLMs) are capable of metabolizing a wide range of OP pesticides and CWNA surrogates as determined using targeted LC-MS/MS and NMR-based metabolomics approaches ^27^. However, HLMs represent a complex enzymatic mixture containing multiple CYP isoforms, esterases, and amidases, making it difficult to identify the specific enzymes responsible for the metabolism of individual OP compounds ^28^. Given the abundance and catalytic promiscuity and previous evidence on OP metabolism by CYP3A4, it is a strong candidate for mediating OP biotransformation in the human liver.

In the present study, we investigated the ability of recombinant human CYP3A4 to metabolize a structurally diverse panel of OP pesticides and CWNA surrogates using targeted LC-MS/MS analysis. We observed that CYP3A4 preferentially metabolizes thion-containing OP compounds relative to their oxon counterparts, consistent with its established role in oxidative desulfuration reactions ^29^. Selected oxon-based CWNA surrogates, including CMP (a cyclosarin analog) and PiMP (a soman analog), also underwent detectable metabolism by CYP3A4 (see Figures S1 and S2 for chemical structures). In parallel, time-dependent inhibition studies revealed that many OP compounds reduced CYP3A4 catalytic activity in both a concentration- and time-dependent manner, suggesting the potential for mechanism-based or quasi-irreversible inhibition or production of a more inhibitory metabolite. In silico docking analysis with 3NXU pdb crystal structure of CYP3A4 revelated that CMP, cyclosarin and sarin as well PiMP and tabun docked at similar binding site in 3D pose. Initial attempts to model CYP3A4 activity using multivariate dimensionality reduction approaches, including principal component analysis (PCA), did not yield robust clustering or predictive separation of OP compounds, likely reflecting the high structural diversity of the dataset and the unusually large/flexible substrate-binding cavity of CYP3A4 ^20,30^. Further, correlation analysis between 10 µM 30 minutes inhibition and molecular weight yielded significant correlation (Pearson, R= - 0.69, P<0.0001). These results overall point to the future development of AI/ML models for predicting CYP3A4 metabolism of OPs as well as authentic CWNA agents.

## 2. Materials and Methods

### 2.1. Chemicals

All OP pesticides used in this study were obtained as neat compounds and were utilized without additional purification. OP pesticides were purchased and surrogates were synthesized as described previously ^31^. CYP3cide (CAS: 526-08-9), Glucose-6-phosphate dehydrogenase (CAS: 9001-40-5), glucose-6-phosphate (CAS: 3671-99-6), NADP+ (CAS: 53-59-8), NADPH (CAS: 53-59-8), and MgCl_2_ (CAS: 7791-18-6) were purchased from Sigma-Aldrich and were HPLC grade. The chemical structures of all OP compounds examined in this work, presented in either oxon or thion form, are shown in Figures S1 and S2 (Supplementary Figures). Compounds containing geometric (E/Z) or stereochemical (R/S) isomers were evaluated as mixed isomeric preparations. Key physicochemical and toxicological properties of the OP compounds—including hydrogen bond donor and acceptor counts, molecular weight, logP, polar surface area, number of rotatable bonds, and reported LD values—are summarized in Supplementary Table S1 ^32–40^.

### 2.2. pFluor50 CYP3A4 Inhibition Assay

CypExpress™ 3A4 (Sigma, MTOXCE3A4), containing full length human recombinant CYP3A4, recombinant human P450 NADPH oxidoreductase, and a proprietary mix of MgCl_2_, magnesium chloride (M9272) and β-nicotinamide adenine dinucleotide phosphate hydrate (N5755), glucose-6-phosphate dehydrogenase (G6PDH;G8529-200UN), glucose-6-phosphate (G6P; G7250), reduced nicotinamide adenine dinucleotide phosphate (NADPH tetrasodium salt; 481973-Millipore) were purchased from Sigma Aldrich (St. Louis, MO). The fluorogenic substrate benzyl 2-(6-(benzyloxy)-3-oxo-3H-xanthen-9-yl) benzoate or dibenzyl fluorescein (DBF) for CYP3A4 (# D0282) was purchased from Chemodex (St. Gallen, Switzerland). Perkin Elmer (Waltham, MA) OptiPlate-384 Black, Black Opaque 384-well Microplates (part no. 6007270) were used for all fluorogenic experiments.

MilliQ water (18.2 MΩ) was used to prepare 0.1 M potassium phosphate buffer (pH 7.4) with potassium phosphate dibasic (0.06958 M) and potassium phosphate monobasic (0.03042 M). OP compounds were diluted in 0.5% DMSO and the entire protocol for running the inhibition (at 10 and 30 minute) for two OP concentrations (1 and 10 µM) was used as described previously in pFlour50 for CYP3A4 ^41^. The rate of increase of relative fluorescence units (RFU) per minute was examined as a measure of the turnover of DBF (Ex 485, Em 535) using a Biotek Synergy H1 plate reader with the slope measured from 15-35 minutes run time and the plate reader parameters were used described previously. CYP3cide was used as positive control for inhibition of CYP3A4 ^41^. Slope normalization was performed as the change in relative fluorescent units per unit time (RFU/time) and data normalization to the average slope of a vehicle control and % inhibition was calculated using Equation 1.

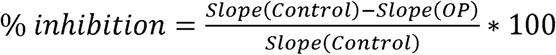

Prism 10.4.2 software (GraphPad software Inc.) was used for data analysis with each inhibition experiment being performed with at least three technical replicates. Mann-Whitney test (Holm-Šídák method of multiple comparisons) was used for pairwise-comparisons with a value of 0.05 for significance.

### 2.3. LC-MS/MS Metabolism Assays

#### CYP3A4 Clearance

BioDuro (Shanghai, China) performed the CYP3A4 LC-MS/MS metabolism study. OP compounds (1 µM final reaction) were pre-incubated with recombinant 15 pmol CYP3A4 at 37 °C for 5 minutes before addition of 1 mM NADPH. 10% of the reaction was separated and subsequently quenched with a 5/10 ng/ml terfenadine/tolbutamide solution in acetonitrile at 0, 5, 30, 60 and 120 minutes. The 4 surrogates were treated in a similar way except the quenching was performed at 0, 5, 15, 30 and 60 minutes. After quenching, the samples were centrifuged (4000 rpm/4°C/15 minutes) followed by mixing the supernatant 1:1 (v/v) with water for LC-MS/MS analysis on either a Q Trap 4500 or API 4000. Midazolam was used as a positive control CYP3A4 substrate. Parameters were calculated as follows:

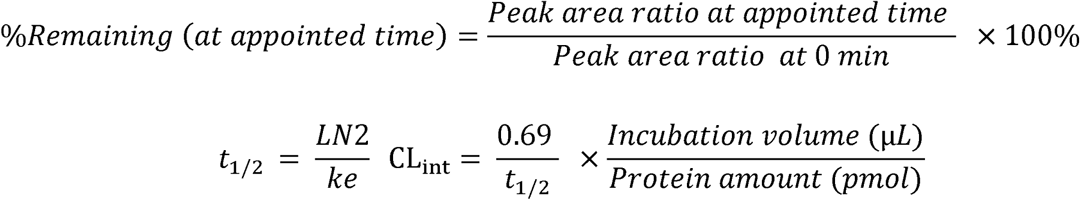

### 2.4. Molecular Docking Analysis

The binding site used for the molecular docking simulations was obtained from the crystal structure of human cytochrome P450 CYP3A4 with bound ritonavir (PDB 3NXU) ^42^. The protein and OP ligand structures were prepared using the same procedure described in our previous work ^31^. Autodock Vina version 1.2.7 was used for molecular docking calculations ^43,44^. The grid box was centered around the iron HEME at 32.67, –21.01, and 21.80 Å and expanded by 20, 18, and 20 points in the x-, y-, and z-directions, respectively, to encompass the nearby residues in the active site. Each OP was docked into the rigid receptor at an exhaustive level of 200, generating up to 20 unique poses.

### 2.5. Correlation and Regression Analyses

Docking-derived binding energies, reactive distances between the phosphorus center and the heme iron, and molecular descriptors were correlated with experimental inhibition and metabolism data. Docking parameter values are presented as average for E/Z or R/S configurations for those pesticides and surrogates wherever applicable. Correlation analyses were also performed between the molecular parameters to investigate their correlation. Individual linear regression and multiple linear regression analyses were performed. Multiple linear regression analysis, although it yielded acceptable R values, showed evidence of data overfitting, so individual linear regression models (Pearson or spearman correlation based on R and P values appropriately) were used for determining correlation. Difference in inhibition at 1 and 10 µM between 10 and 30 minutes as well as individual values were used in the correlation analyses along with LC-MS/MS detection of the percent of OP compound remaining between 0 and 1 or 0 and 2 hours correlated with the molecular parameters, docking-derived binding energies, or reactive distances. Correlation data is presented as the correlation coefficients (R) and significance values (P) of the regression line with the corresponding equation as calculated using GraphPad Prism. Throughout the paper, Pearson or Spearman correlations have been used which were selected based on R and P values.

## Results

### CYP3A4 is inhibited by oxons and thions in a time-dependent manner

CYP3A4 activity was quantified using the previously developed pFluor50 fluorogenic method with DBF as a substrate ^41^. In this study, the same protocol was used to investigate the activation or inhibition of 47 OP compounds on CYP3A4 activity. CYP3cide was used as a positive control for CYP3A4 inhibition (Figure 1 A-B). We found that 14 thions at 1 µM and 19 at 10 µM showed time-dependent variation in CYP3A4 inhibition between 10- and 30-minutes incubation time (Figure 1A). While 14 out of 26 oxons at 1 µM and 18 at 10 µM showed significant variation in CYP3A4 inhibition between 10 and 30 minutes (Figure 1B). The overall summary of the data at 1 and 10 µM between 10 and 30 minutes is presented in a heatmap (Figure 1C). Many OP compounds, such as PPE, PMT, MTN, PM and PiMP at 1 µM and IPP, IFP, NEDPA and MMP at 10 µM, activated CYP3A4 at 10 minutes but inhibited at 30 minutes Furthermore, some OP compounds, such as DMA, FMP, EPP, EMP and DMX, inhibited CYP3A4 at 10 minutes but activated it at 30 minutes. These time-dependent differences in CYP3A4 suggest either a time-dependent change in interaction with CYP3A4 (mechanism-based inhibition) or the formation of a secondary metabolite with likely different inhibitory properties.

**Figure 1.**
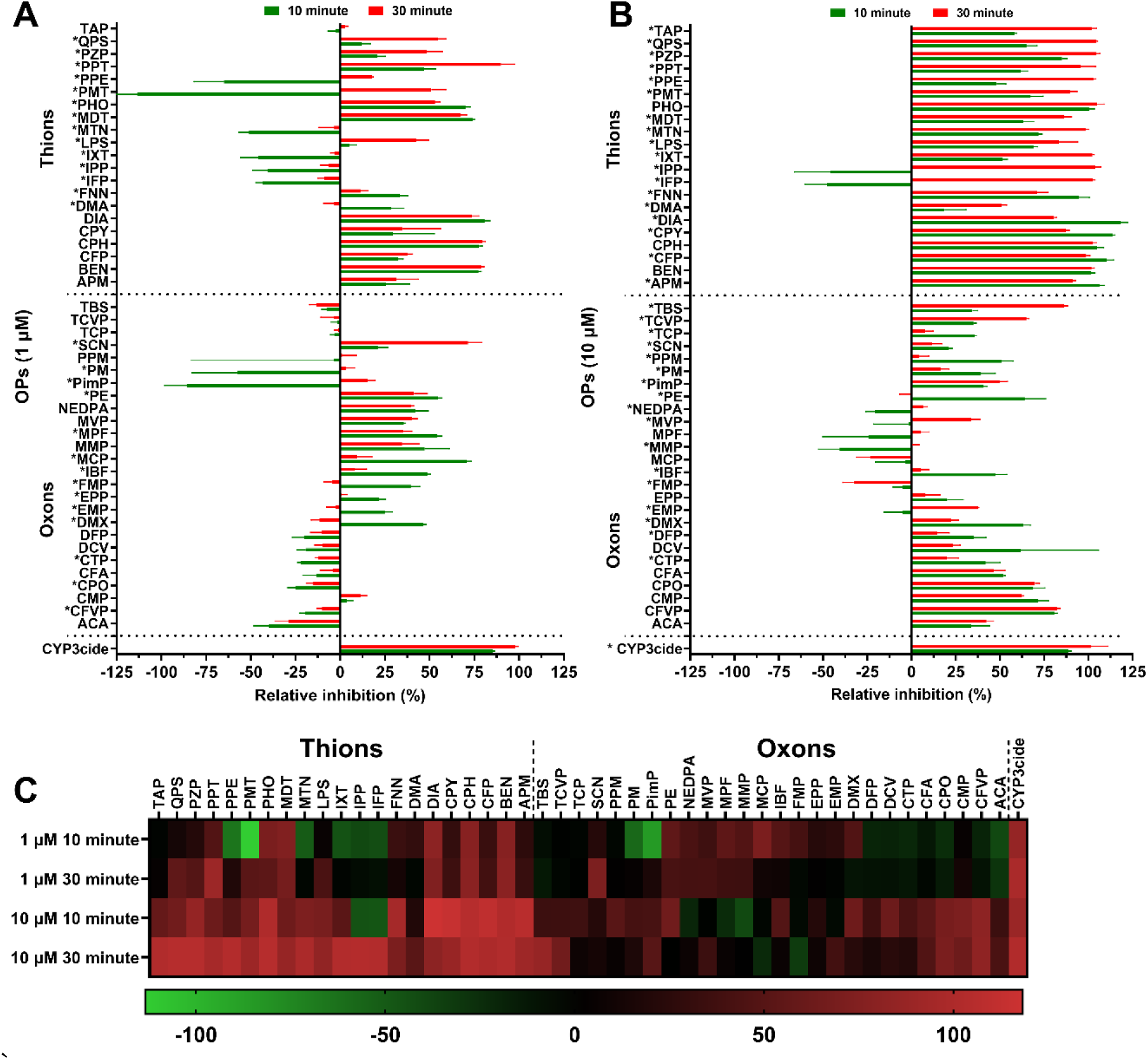
OP-mediated relative inhibition using pFluor50 fluorogenic assay for CYP3A4 activity. A. Relative inhibition of CYP3A4 activity using pFluor50 fluorogenic method with DBF as a substrate ^41^ by thions and oxons at 1 µM concentration. B. Relative inhibition of CYP3A4 activity by thions and oxons at 10 µM concentration. C. Heatmap showing variations in CYP3A4 activity due to OP inhibition at 1 and 10 µM between 10 and 30 min. * Represents OP compounds wherein the inhibition of CYP3A4 changes significantly between 10 and 30 min. CYP3cide was used as positive control for inhibition of CYP3A4.

### Thions are selectively more sensitive to CYP3A4 as compared to oxons

LC-MS/MS experiments were also performed to detect the clearance of OP compounds after incubation with CYP3A4 (Table 1). The amount of each OP compound was quantified at different times up to 1 hour (CWNA surrogates) or 2 hours (all other compounds tested). Midazolam was used as a positive control and was metabolized completely within 30 minutes of incubation. We found that all 12 thions were degraded by more than 50% within the timeframe tested, and IPP, PHO and FMN were metabolized completely within 5 minutes of incubation. TEM, MDT and MTN were metabolized the slowest, showing a reduction of around 50% in 2 hours. Further QPS, IXT, TAP, DIA, PMT, PPT and QPS were 75% metabolized after 2 hours (Figure 2B). Among the oxons, only FMP and EPP were degraded by more than 80% after 2 hours, while other OP compounds were metabolized less than 50%. ACA, PFN, MVP and DCV were not degraded at all, while TCVP, PPM, CFVP and TCP were metabolized by around 50% after 2 hours (Figure 2A). CMP (cyclosarin surrogate) and PiMP (soman surrogate) were metabolized more than 80%, while EMP (a VX surrogate) and NEDPA (a tabun surrogate) were not metabolized at all during the 1-hour exposure, suggesting that CYP3A4 was selective towards specific OP surrogates (Figure 2C). Overall, thions (mean 23.33%) were metabolized more than twice as fast as oxons (mean 68.82%) after 1 hour of CYP3A4 incubation with significant (P=0.0015) mean difference (–45.49 ± 12.74) and with a 95% confidence interval (–71.79 to –19.19) (Figure 2D).

**Table 1.**
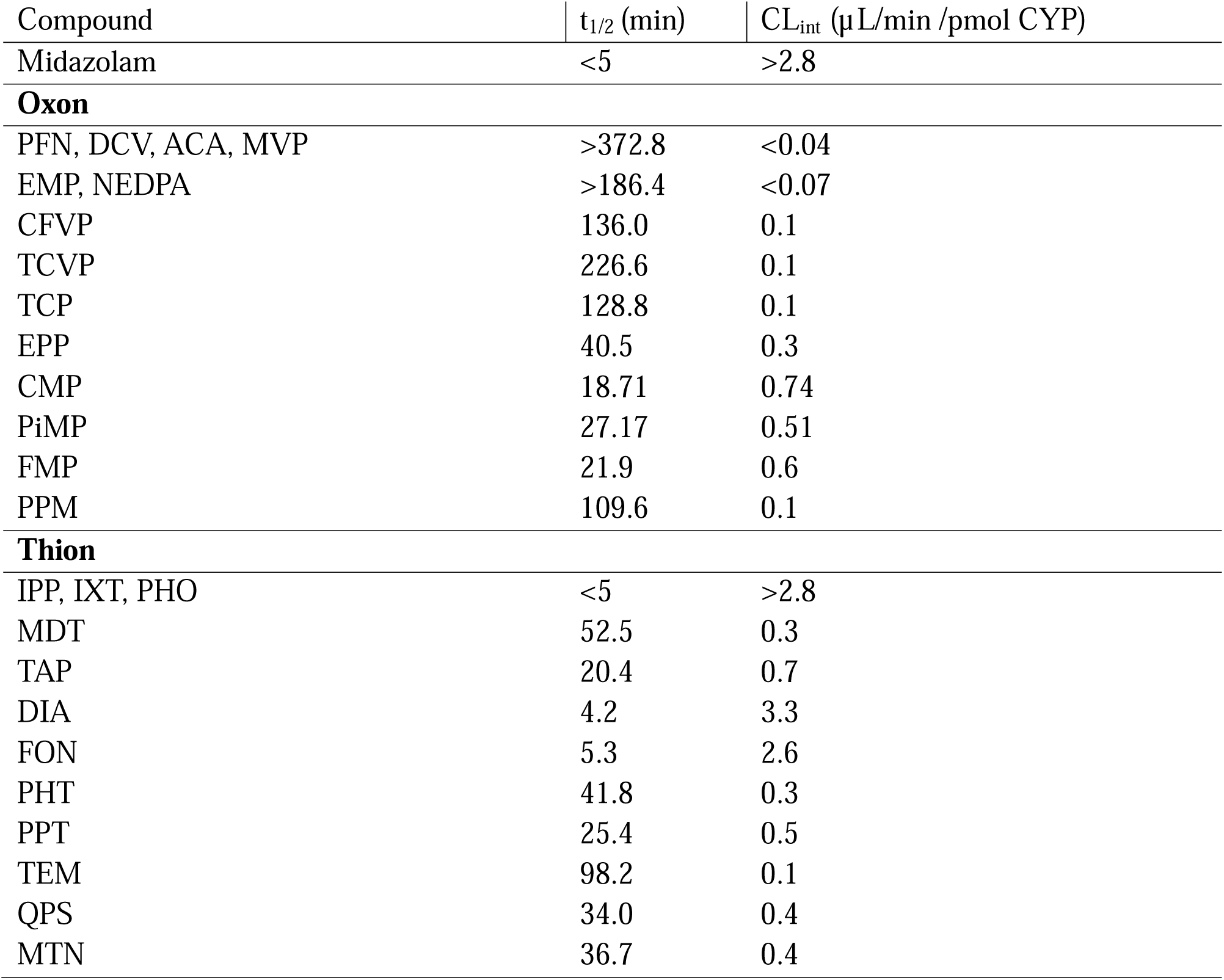
Half-life and intrinsic clearance of OP pesticides by recombinant CYP3A4 as measured by the disappearance of the parent OP derivative.

**Figure 2.**
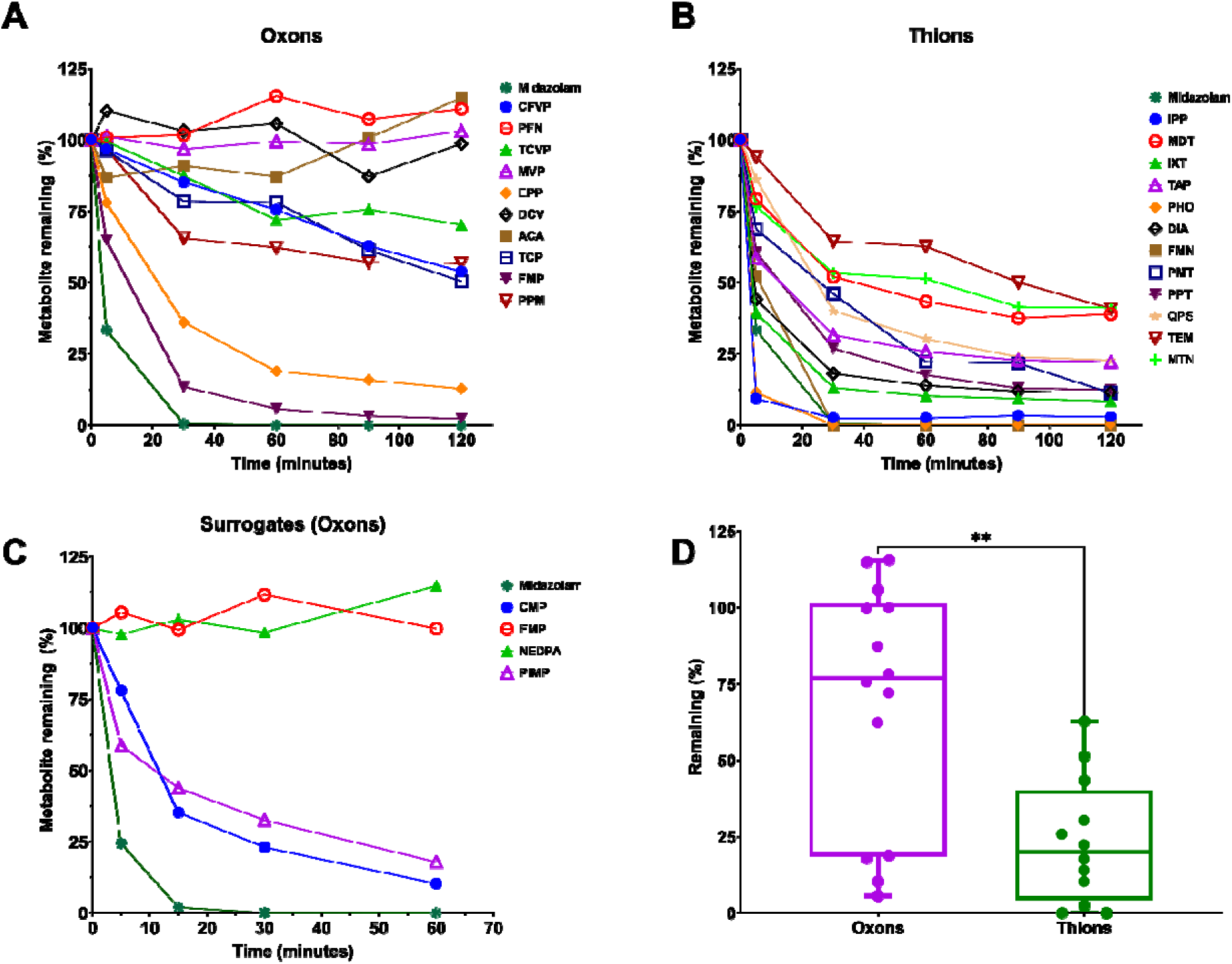
Metabolism of OP compounds by CYP3A4 measured by disappearance of the parent. **A**. LC-MS/MS metabolism of oxons by recombinant CYP3A4 after 2 hrs. **B**. LC-MS/MS degradation of thions by recombinant CYP3A4 after 2 hrs. **C**. LC-MS/MS metabolism of CWNA surrogates by recombinant CYP3A4 after 1 hr. **D**. CYP3A4 mediated metabolism of thions was significantly higher as compared to oxons based on mean averages after 1 hr. **P<0.01. Purple circles represent oxons, green circles thions,

### Molecular Docking analysis shows correlation between the individual docking parameters and fluorogenic assay-based inhibition or LC-MS/MS-based degradation experiments

Autodock Vina was used to perform rigid docking the in the CYP3A4 enzyme (PDB ID : 3NXU) with each of the oxons (including CWNA surrogates) or thions in R/S or E/Z configurations wherever applicable. (Supplementary Figure S1 and S2). 3NXU has one of the highest resolutions and has been used previously for CYP3A4 docking ^20,42,45,46^ as it contains Ritonavir bound to the heme center. We previously presented a correlation analysis between fluorogenic assay-based inhibition and LC-MS/MS metabolism experiments and the docking parameters for OP compounds in CYP2C9 ^31^. Here, we also included correlation with various molecular parameters for each OP including molecular weight (MW), predicted or measured LD_50_, hydrogen-bond acceptors (HBA), hydrogen-bond donors (HBD), polar surface area (PSA), number of rotatable bonds, octanol-water partition coefficients (logP), and van der Waals radii (VWR) (Supplementary Table 1) along with the docking parameters (binding energy (BE), reactive distance (D), and binding efficiency (BE_eff_) which is docking score divided by molecular weight of the compound) for the best scored pose and the pose with the shortest reactive distance (SD, BE(SD), BE_eff_(SD)) (Supplementary Table 2). We determined the correlation between each of the individual parameters (Supplementary Figure S3) to investigate which parameters might be used together when performing multiple variable regression and principal component analysis and observed that MW was significantly correlated with logP, rotatable bonds, and VWR while HBA was correlated with PSA, HBD was correlated with logP, logP was correlated with PSA, and PSA was correlated with VWR (Supplementary Table 11).

We first analyzed the correlation of each parameter individually with the time-dependent difference in inhibition at 1 µM between 10 and 30 minutes and found that MW, VWR, BE and BE(SD) were significantly correlated (Supplementary Table 3), while MW and BE_eff_ were significantly correlated to the time-dependent difference in inhibition at 10 µM between 10 and 30 minutes (Supplementary Table 4). We then investigated the correlation of each individual parameter with the individual measurements of inhibition at 1 and 10 µM at the 10- and 30-minute timepoints. At the 1 µM concentration, none of the parameters were significantly correlated with the individual inhibition measurements for any of the determined conditions at either the 10- or 30-minute timepoints (Supplementary Table 5-6). We found that at the 10 µM concentration and the 30-minute timepoint, there was a positive correlation between inhibition and the MW of the OP compounds (R = 0.69; P <0.0001), while there was a negative correlation between inhibition and the BE at the shortest reactive distance (R = –0.48; P<0.001) (Figure 3A-B). Furthermore, all of the thions with the exception of FNN, PHT and DMA exhibited nearly 100% inhibition at the 30-minute timepoint, suggesting either mechanism-based inhibition with the thions in CYP3A4 is occurring by 30 minutes or that the formation of secondary metabolites after thion degradation is occurring by 30 minutes and the secondary metabolites are very potent inhibitors of CYP3A4. Oxons, on the other hand, showed greater variation in inhibition of CYP3A4 at the 30-minute timepoint (Figure 3A-B). The logP (R = 0.57; P<0.0001), VWR (R = 0.60, P<0.0001), and BE (R = –0.48, P< 0.001) parameters were also significantly correlated (Figure 3C-E) with inhibition at the 30-minute timepoint. Rotatable bonds (R = 0.33, P<0.05), BE_eff_ (R = 0.29, P<0.05) and BE_eff_(SD) (R = 0.31, P<0.05), were also significantly correlated with inhibition, but with more modest R values (Supplementary Table 7). Furthermore, inhibition at 10 µM at the 10-minute timepoint also showed significant correlation with BE(SD) (R = – 0.41; P<0.01), MW (R = 0.43; P<0.01), HBD (R = –0.36; P<0.05), logP (R = 0.35; P<0.05), VWR (R = 0.40; P<0.01) and BE (R = –0.45; P<0.01) (Figure 3F, Supplementary Table 8). Although inhibition at the individual 10 µM concentration at the two different timepoints showed modest but significant correlation with selected molecular parameters, there was no significant correlation between the parameters and the time-dependent difference in inhibition at either concentration (Supplementary Figure S3).

**Figure 3.**
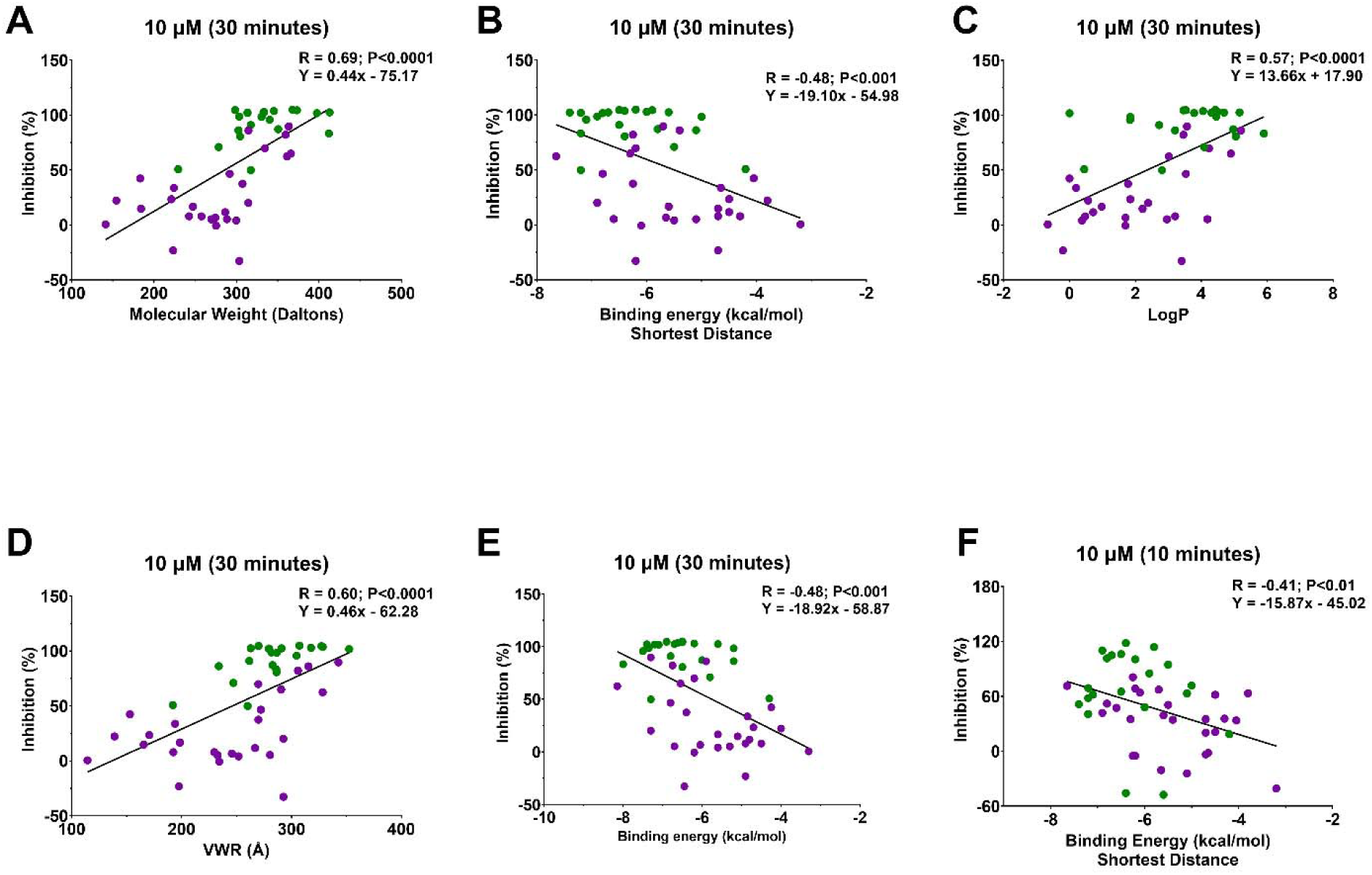
Correlation analysis between molecular parameters and pFluor50-based CYP3A4 inhibition at 10 µM. **A-E**. Correlation analysis between 10 µM 30-minute inhibition with MW (Spearman Rho R = 0.69; P<0.0001), BE(SD) (Spearman Rho R = –0.48; P<0.001), logP (Spearman Rho R = 0.57, P<0.001), VWR (Spearman Rho R = 0.60; P<0.0001), BE (Spearman Rho R = –0.48; P<0.001). B. Correlation analysis between 10 µM 10-minute inhibition with BE(SD) (Spearman Rho R = –0.41; P<0.01). Purple dots represent oxons, green dot thions.

We then investigated the correlation between the molecular parameters and the LC-MS/MS detected metabolism of OP compounds by CYP3A4 for 1 or 2 hours. The concentration of parent OP remaining after 2 hours (% remaining) was significantly correlated with BE(SD) (Figure 4A; R = 0.58; P<0.01). With the exception of FON and EMP (which had weaker BE(SD) and high metabolism), these data seemed to cluster into three sets. The first set (Figure 4A, red circle) had stronger BE(SD) but high metabolism and included mostly thions such as IPP, DIA and others with only one oxon, FMP; the second set (Figure 4A, purple circle) had weaker BE(SD) and the lowest degradation and included only oxons such as DCV, ACA, PFN and MVP; and the third set which was intermediate for both of the parameters (Figure 4A, orange circle) included the thions MTN, MDT and TEM and the oxons TCP, TCVP, CFVP and PPM. Stronger binding refers to more negative BE while weaker BE refers to less negative BE. Further, the BE (R = 0.57; P<0.01), VWR (R = -0.57; P<0.01), MW (R = –0.46; P<0.05), logP (R = –0.45; P<0.05) and rotatable bonds (R = –0.48; P<0.05) were also significantly correlated with the LC-MS/MS detected metabolism after 2 hours (Figure 4B, Supplementary Table 9). Additionally, the BE (R = 0.46; P<0.05), VWR (R = –0.45; P<0.05) and logP (R = -0.44; P<0.05) were also correlated with the percent of OP remaining (%) after 1 hour of CYP3A4 metabolism (Figure 4C-D, Supplementary Table 10) while other parameters did not show any significant correlation. We also performed a correlation analysis between fluorogenic-detected inhibition and LC-MS/MS-detected metabolism. Inhibition at the 10 µM concentration and 30-minute timepoint and the percent of OP remaining after 2 hours (%) showed a negative correlation for thions (Figure 4E; R = 0.70, P<0.05) as well as for all OPs (Figure 4F; R = -0.46, P<0.05). Additionally, the percent of oxon OP remaining after 2 hours and the percent of all OPs (oxons and thions) remaining after 1 hour also showed significant correlation between the two experimental datasets (Supplementary Figure S4A-D). Although we also performed a PCA analysis for all of the parameters, as well as multiple linear regression analysis with selected parameters, there were no significant results in this analysis (data not shown). Thus, our correlation analysis for CYP3A4 was confined to the correlation between individual parameters and experimental data.

**Figure 4.**
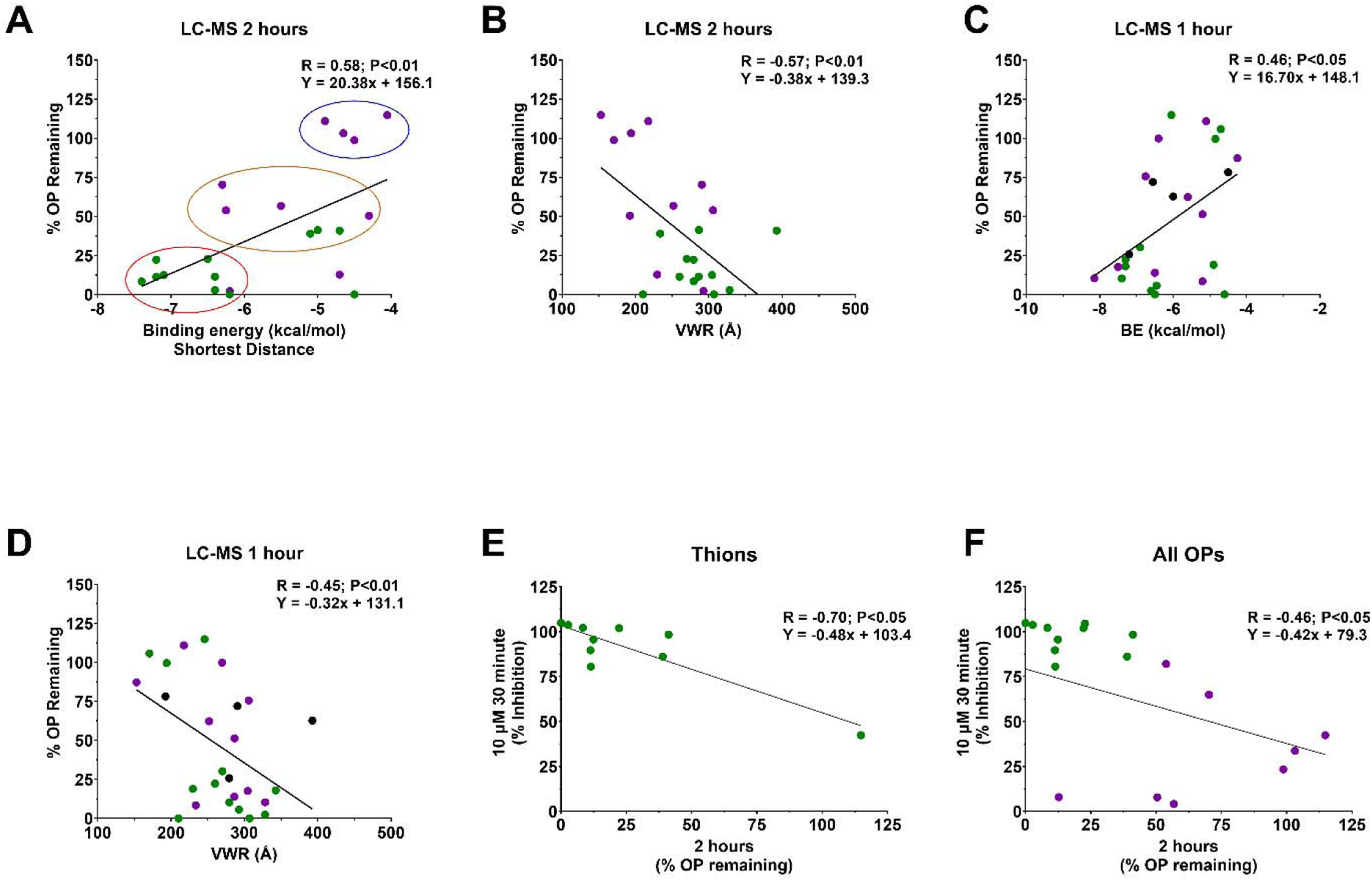
Correlation analysis between molecular parameters and LC-MS/MS degradation. **A.** Correlation analysis between percentage of parent remaining in the LC-MS/MS metabolism experiments after 2 hrs and BE(SD) (R = 0.58; P<0.01) and VWR (R = –0.57; P<0.01). **B**. Correlation analysis between percentage of parent remaining in the LC-MS/MS metabolism experiments after 1 hr and BE (R = 0.46; P<0.05) and VWR (R = –0.45; P<0.01). **C**. Correlation analysis between 10 µM 30 minutes and % parent OP remaining after 2 hrs for thions (R = – 0.70; P<0.05) and all OP compounds (R = –0.46; P<0.05). Purple dots represent oxons, green sot thions. All correlations in this figure are Pearson or Spearman based on R and P values.

We also explored the interactions between CYP3A4 and the CWNA surrogates compared with the authentic CWNAs for the docked pose with the shortest reactive distance. 3D docked poses of CMP, cyclosarin and sarin for both R and S conformations were found to bind in a similar site and orientation in the enzyme bindingactive site (Figure 5A-B). Cyclosarin and CMP had close proximity with six identical residues in addition to the heme - namely HEM442, Ile301, Ser119, Leu482, Leu211, Ala305, Phe 304 demonstrating that they bind within the same region (Figure S5A-B). 3D docked poses for PiMP and soman were also found to bind at a similar site and orientation active site (Figure 5C-D). PiMP and soman had close proximity to HEM442, Ile301, Ala305, Phe 304, Ile369 and Ser119 with both having a CH-π interaction between the aromatic HEM442 and the methyl group bonded to the phosphorus (Figure S5C-D). 3D docked poses of EMP and VX, although bound at a similar site, were in different orientations (Figure S6A-B). EMP and VX had close proximity with HEM442, Thr309, Ser119, Phe304, Ile301, Ala305, and Arg105 (Figure S5E-F). 3D docked poses of NEDPA and tabun were in a similar orientation (Figure S6C-D). NEDPA and tabun had close proximity with HEM442, Thr309, Ser119, Phe304, Ile301, Ile369, and Ala305 (Figure S5G-H). The common residues in the binding pocket between the surrogates and authentic CWNAs suggest that the metabolism of the surrogates may also be predictive of CYP3A4 metabolism of authentic CWNAs.

**Figure 5.**
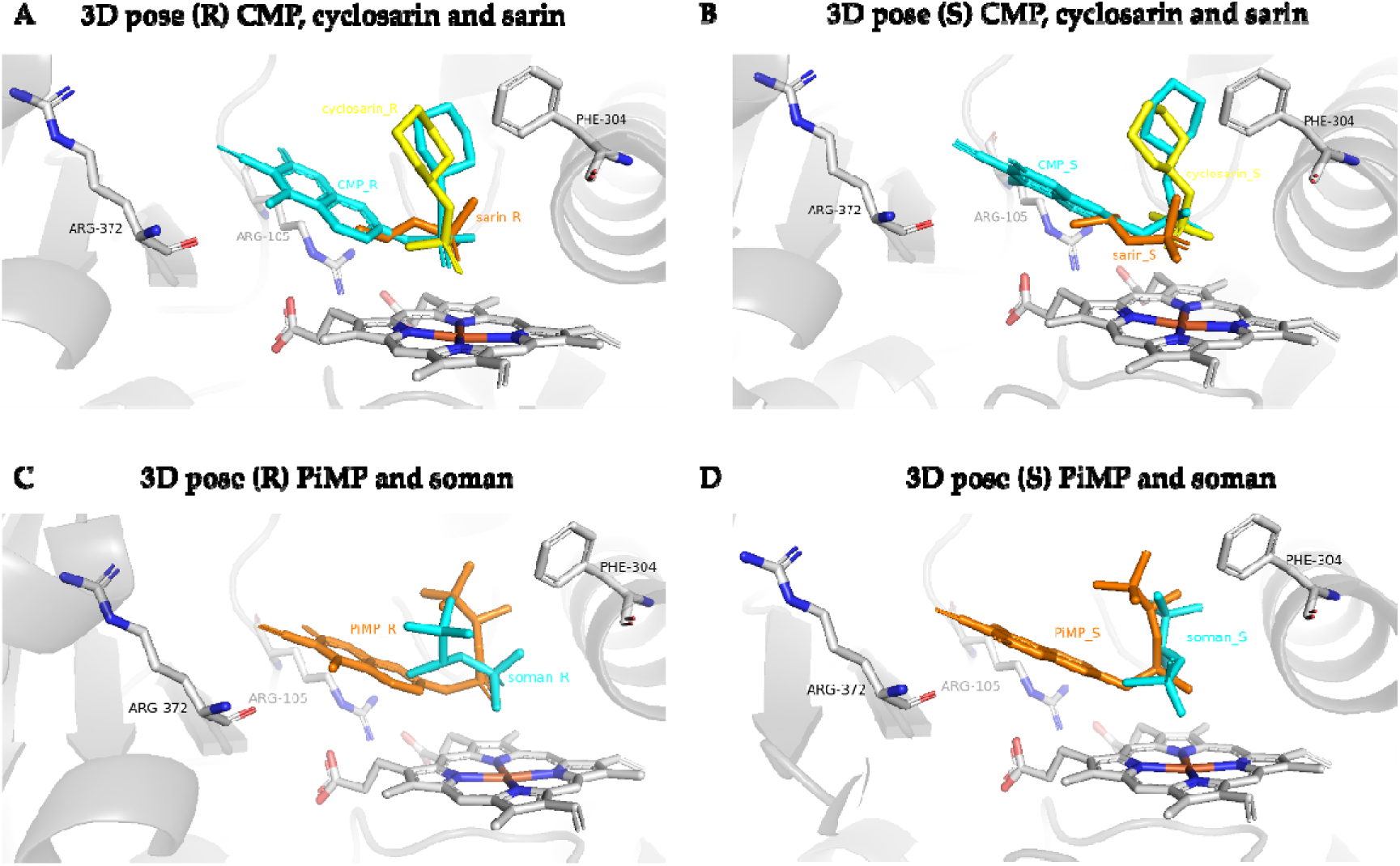
3D docking poses (shortest reactive distance) for the R and S enantiomers of CMP/cyclosarin/sarin and PiMP/soman with 3NXU PDB structure of CYP3A4. **A**. 3D docking poses for CMP (*R*), cyclosarin (*R*) and sarin (*R*). **B**. 3D docking poses for CMP (*S*), cyclosarin (*S*) and sarin (*S*). **C**. 3D docking poses for PiMP (R) and Soman (R). **D**. 3D docking poses for PiMP (S) and Soman (S). Cyclosarin is yellow, sarin and CMP is cyan in color in panel A-B while PiMP is orange and soman is cyan in panel C-D.

## Discussion

OP poisoning is a major health concern throughout the world due to the alarmingly high rate of OP use in suicide attempts, with 15% of total (150,000-200,000) suicidal deaths worldwide being from OP poisoning ^2,3,16^. Additionally, residual OP can be detected in most inorganically grown crops, further contributing to chronic exposure for both consumers of such fruits and vegetables as well as the agriculture workers who grow and harvest them ^7–9^. The current therapies targeting AChE remain inadequate particularly in the CNS, so new therapies for OP poisoning are urgently needed ^18^.

We previously developed pFluor50, a fluorogenic assay for CYP3A4, and used this assay to determine the kinetic activity of the enzyme using DBF as a substrate. We further validated this assay using CYP3cide as inhibitor of CYP3A4 ^41^ demonstrating an increase in inhibition with both time and dose. Here we have used this assay to screen a number of OPs for inhibition/activation of CYP3A4 at two different timepoints (10 and 30 minutes) and two different dose levels (1 and 10 µM). Many OPs such as PPE, PMT, PPM, PM and PiMP were activators at 10 minutes but inhibitors at 30 minutes. FMP was the only OP wherein the activation increased from 10 to 30 minutes, while DIA, CPY and APM showed a decrease in inhibition from 10 to 30 minutes (Figure 1A-B). The marked increase in inhibition from 10 to 30 minutes is consistent with a time-dependent inhibitory component, which may involve mechanism-based inactivation or formation of inhibitory metabolites, although additional studies would be required to distinguish between these possibilities. The hypothesis of formation of secondary metabolites with variations in inhibition is consistent with previously reported lapatinib metabolism by CYP3A4 and CYP3A5 ^24,47–49^. The majority of thions, such as APM, BEN, CPH, IXT and IPP, at 10 µM and 30 minutes were complete inhibitors (showing ∼100% inhibition). Indeed, among the thions only FNN, PHT and DMA showed less than 80% inhibition at 10 µM and 30 minutes. In contrast, among the oxons, only TBS, PiMP and CFVP, showed an inhibition of greater than 80% under the same conditions. These data further highlight the differences in activity between of these two classes of OPs (Figure 1B). These findings suggest that future studies focusing on metabolite identification via LC-MS/MS followed by the determination of inhibition of CYP3A4 by the identified individual metabolites would be of interest.

Previous research, including ours, has focused on using human liver microsomes (HLMs) to investigate the metabolism of OPs using NMR as well as LC-MS/MS (Sams, Mason and Rawbone, 2000; Buratti *et al.*, 2003; Rose *et al.*, 2005; Ellison *et al.*, 2012; Agarwal *et al.*, 2023). However, due to the presence of a large number of candidate enzymes in HLMs it is difficult to predict which enzyme(s) are responsible for any OP metabolism that is detected using such techniques ^54^. To gain additional insights, we previously used CYP2C9 to study OP metabolism and found a number of different OPs that were metabolized ^31^. However, there were a number of OPs that were not degraded by CYP2C9, so herein we investigated which Ops CYP3A4 might metabolize. We found that, in general, thions were metabolized more quickly than oxons overall after 1 hour (Figure 2D). This combined with the inhibition data at the 10 µM concentration and 30-minute timepoint suggests that CYP3A4 may have higher binding affinity for thions (or the metabolites thereof). Previously, other thions such as chlorpyrifos ^55^, diazinon ^26^, dimethoate ^56^, parathion ^23^, methyl parathion (Ellison *et al.*, 2012), malathion ^58^, azinphos-methyl and phosmet ^59^ were shown to be desulfurized to their oxon form by CYP3A4. In this study, a number of additional OPs, including oxons, were also studied. Our findings confirmed these previous results (as DIA, PMT, and MTN were metabolized in our LC-MS/MS analysis (Figure 2B)) as well as extended the them to additional compounds.

We also performed molecular docking analysis to correlate these experimental results with the predicted binding affinity of the OPs with CYP3A4. In our previous work, we correlated the docking parameters with the experimental data ^31^, but here, in addition to the docking results, we have also examined the correlation of a number of molecular properties with the experimental data. Many of the properties examined demonstrated significant positive or negative correlation with experimental data (Figure 3–4, Supplementary table 3-10). Additionally, the comparison of the BE(SD) with the metabolism of the OPs after 2 hours (R=0.58, P<0.01) separated the OPs into three distinct sets with the majority of thions metabolized completely while yielding a lower BE(SD) and the oxons with higher BE(SD) demonstrated the lowest metabolism (Figure 4A). Furthermore, the inhibition data at 10 µM and 30 minutes showed modest but significant correlation with many of the properties like MW and logP in addition to the BE(SD). However, the inhibition data at the 1 µM concentration did not show any significant correlation, highlighting that higher concentrations might be required for accurately predicting the CYP3A4 activity (perhaps due to the large active site of CYP3A4 compared to the small size of some of the OP compounds). Finally, the correlation between the inhibition data at 10 µM and 30-minutes showed the strongest correlation with the LC-MS/MS detected metabolism after 2 hours (R = – 0.70, P<0.05). Although the single-parameter correlation models contained herein showed only modest correlation, future work exploring more rigorous methods such as Molecular Mechanics Poisson–Boltzmann Surface Area (MM-PBSA), may provide a more predictive binding affinities.

Importantly, our docking analysis comparing CWNA surrogates with the authentic agents demonstrated that they reside in a similar cavity generated by similar residues in CYP3A4 active side (e.g., with 7 or 8 residues for cyclosarin or soman being the same as those for CMP or PiMP, respectively), points to the possibility of similar interaction with CYP3A4. Indeed, we found that CMP and PiMP were both metabolized by more than 80% after 1 hour of incubation with CYP3A4 (Figure 2C). Collectively, these *in vitro* and *in silico* findings highlight the important role of CYP3A4 in OP metabolism and support the potential of integrating experimental and *in silico* data. Future research using CYP3A4 to metabolize authentic CWNAs would be particularly enlightening.

## Patents

Not applicable.

## Supplementary Materials

Supplementary materials are attached separately.

## Author Contributions

Conceptualization, P.S. and C.A.M.; methodology, P.S., A.M.N., B.T.S.; software, P.S., A.M.N., B.T.S., C.A.M. and C.M.H.; validation, C.A.M., C.M.H., and T.M.; formal analysis, P.S. and C.A.M.; investigation, P.S., A.R., A.M.N., B.T.S.; resources, C.A.M., C.M.H. and S.E.; data curation, P.S.; writing—original draft preparation, P.S.; writing—review and editing, P.S., C.A.M., C.M.H., T.R.L. and S.E; visualization, P.S.; supervision, C.A.M., C.M.H., and S.E.; project administration, C.A.M. and S.E.; funding acquisition, C.A.M. and S.E. All authors have read and agreed to the published version of the manuscript.

## Funding

This research was funded by Defense Threat Reduction Agency, an affiliate of Department of Defense, grant number HDTRA11910020.

## Institutional Review Board Statement

Not applicable.

## Informed Consent Statement

Not applicable.

## Data Availability Statement

Data are contained within the article.

## Acknowledgments

The authors would like to thank Dmitriy Uchenik (Shared Instrumentation Facility Director, College of Pharmacy) instrument facility for use of the BioTek plate reader. Computational resources were provided by the Ohio Supercomputer Center.

## Conflicts of Interest

The authors declare the following competing financial interest(s): S.E. is founder and owner of Collaborations Pharmaceuticals, Inc. and T.R.L. is an employee of Collaborations Pharmaceuticals, Inc.

## Abbrevations

AChE: Acetylcholinesterase
ACA: Acephate
APM: Azinphos-methyl
BE: binding energy
BNS: Bensulide
CFVP: Chlorfenvinphos
CPH: Chlorphoxim
CPY: Chlorpyrifos
CPO: Chlorpyrifos Oxon
CTP: Crotoxyphos
CFA: Crufomate
CFP: Cyanofenphos
CYP: cytochrome p450
CWNA: chemical warfare nerve agent
DFP: Diisopropyl fluorophosphate
DIZ: Diazinon
DCV: Dichlorvos
DEP: Diethylparaoxon
DFP: Diisopropyl fluorophosphate
DMX: Dimefox
DMA: Dimethoate
DMP: Dimethyl paraoxon
EPP: Ethoprophos
Eres: resorufin ethoxy ether
FMP: Fenamiphos
FNN: Fenthion
FON: Formothion
IFP: Iodofenphos
IBF: Iprobenfos
IPP: Isofenphos
IXT: Isoxathion
LPS: Leptophos
MTN: Malathion
MPF: Mephosfolan
MMP: Methamidophos
MDT: Methidathion
MVP: Mevinphos
MCP: Monocrotophos
OP: organophosphorus compound
PHO: Phosalone
PFN: Phosfolan
PHT: Phosmet
PPM: Phosphamidon
PPE: Pirimiphos-ethyl
PZP: Pyrazophos
PPT: Pyridaphenthion
QNP: Quinalphos
RRB: Rotatable bonds
SCN: Schradan
TEM: Temephos
TCVP: Tetrachlorvinphos
TAP: Triazophos
TBS: Tribufos
TCP: Trichlorphon.

## Notes

### Competing Interest Statement

The authors have declared no competing interest.

