## supplementary file for "Organophosphorus pesticide and nerve agent surrogate metabolism by human CYP3A4"

**Supplementary Figures**

**Supplementary Figure 1. Structures of all of the oxon organophosphorus compounds used in the experiments and docking analysis.**

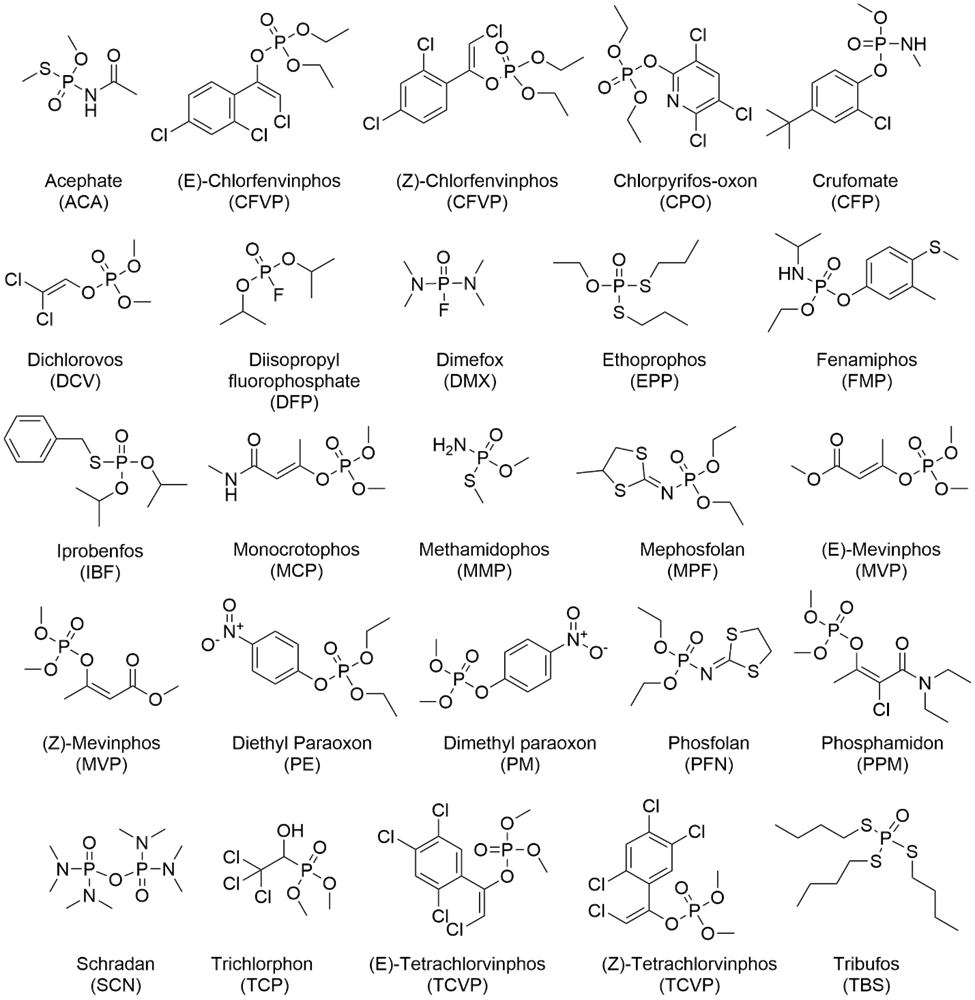

**Supplementary Figure 2. Structures of all of the thion organophosphorus compounds used in the experiments and docking analysis.**

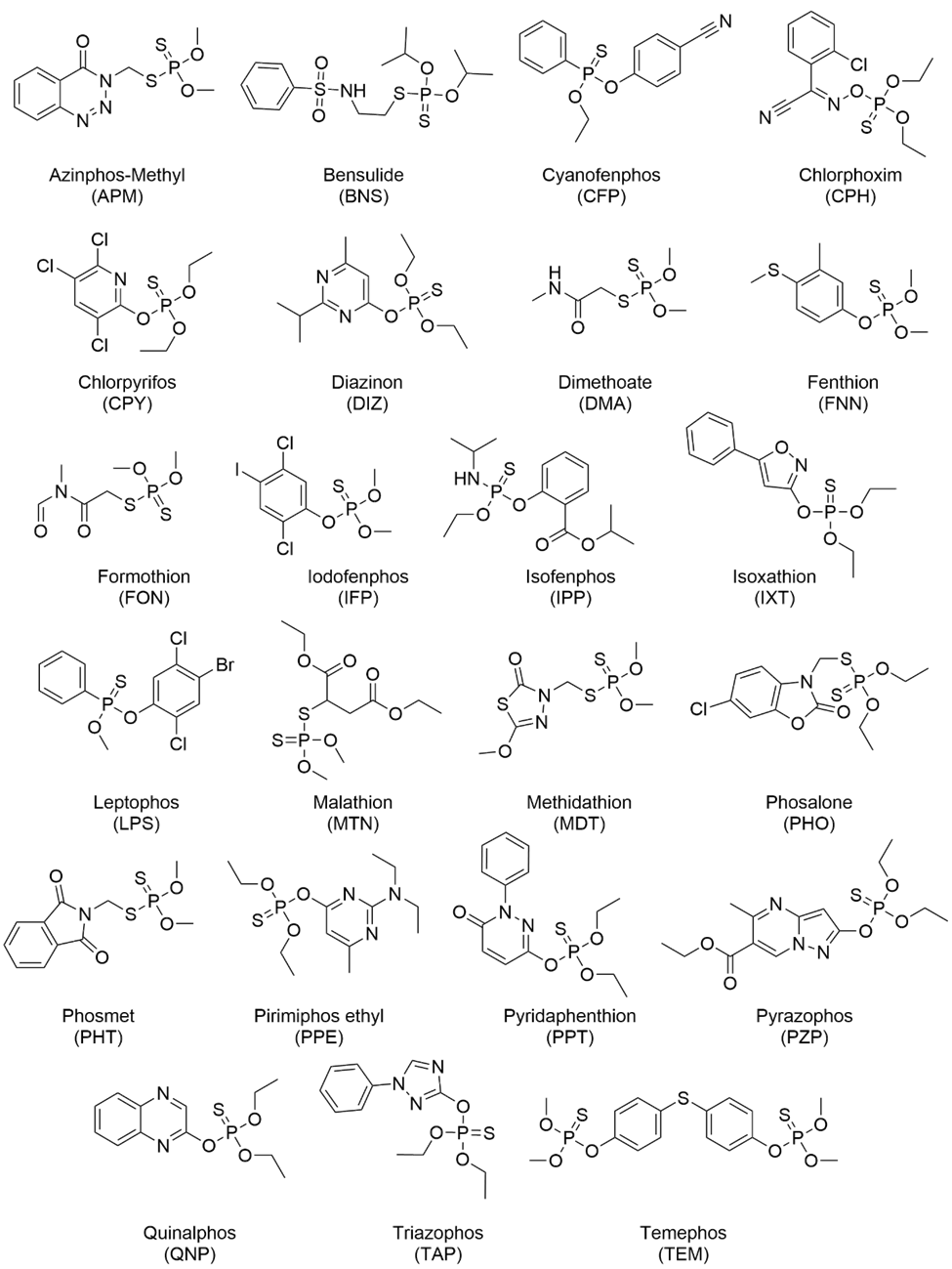

**Supplementary Figure S3. Correlation matrix between docking parameters and molecular parameters for the 47 OPs used in the paper**. Averages were taken for R/S or E/Z pairs of OP. Blue color indicates strong positive correlation while red color indicates strong negative correlation. Pearson correlation has been used. MW – Molecular weight, LD50 – Lethal dose 50%, HBA – hydrogen bond acceptor, HBD – hydrogen bond donor, LogP – Partition coefficient lipophilicity for lipids over water, PSA – polar surface area, VWR – Vander Waal radii, BE – Binding energy, BE_eff_ – Binding efficiency, top pose – best pose based on stronger (more negative) binding energy.

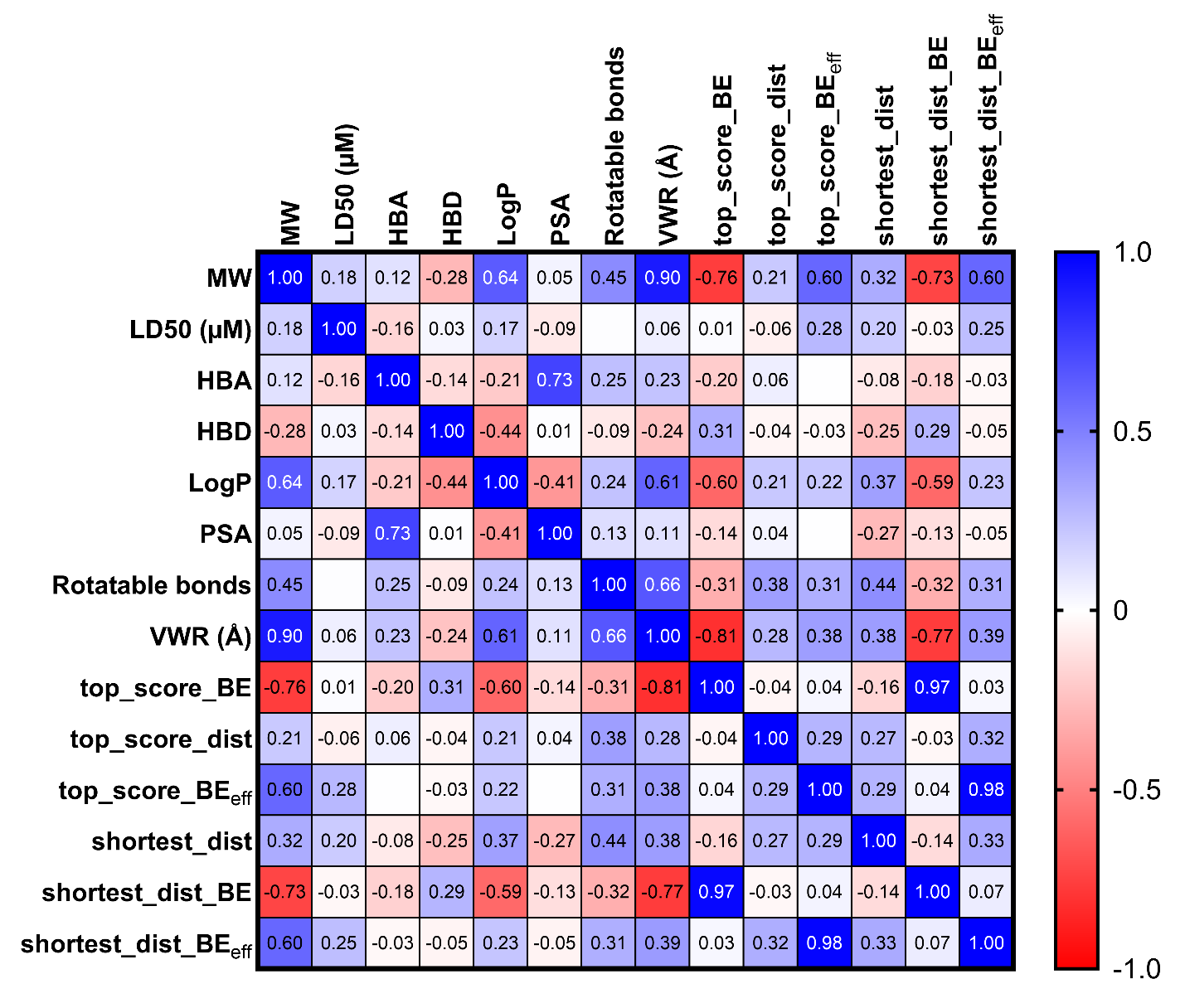

**Supplementary Figure S4. Correlation analysis between OPs (thions in green and oxons in purple) between the CYP3A4 pFluor50-based assay inhibition at 10 µM and 30 minutes and the LC-MS/MS analysis of OP metabolism after 1 or 2 hours. A.** Correlation analysis (R = 0.3, P:NS) between % of oxon remaining after 2 hours and inhibition by oxons at 10 µM and 30 minutes**. B.** Correlation analysis (R = -0.41, P:NS) between % of OP remaining after 1 hour and inhibition by OPs at 10 µM and 30 minutes**. C.** Correlation analysis (R = -0.61, P:NS) between % of thion remaining after 1 hour and inhibition by thions at 10 µM and 30 minutes**. D.** Correlation analysis (R = -0.03, P:NS) between % of OP remaining after 1 hour and inhibition by oxon at 10 µM and 30 minutes**.** NS: Not significant. All correlations are Pearson.

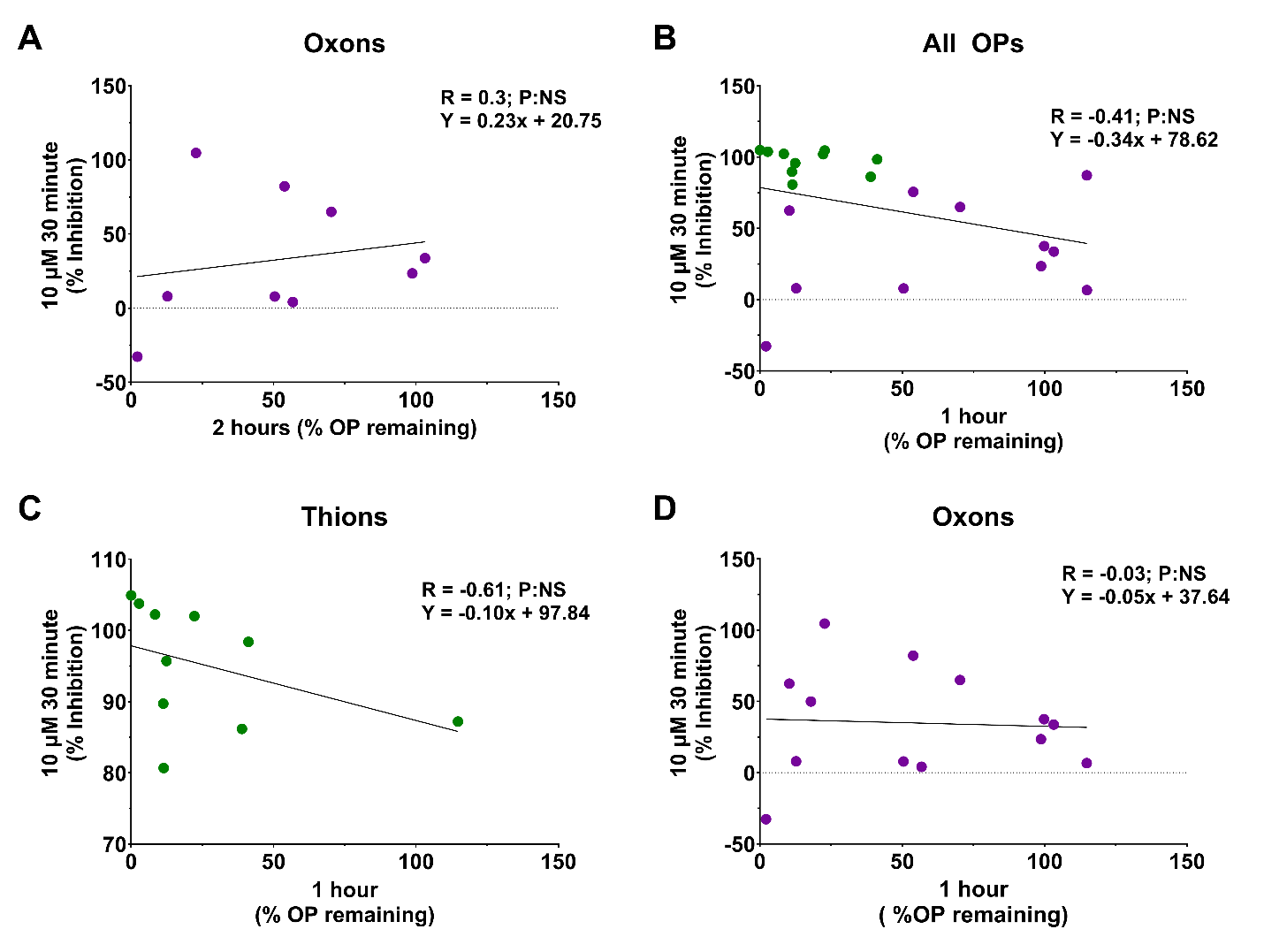

**Supplementary Figure S5. 2D interaction diagrams between CWNA surrogates and CWNAs with the 3NXU PDB structure of CYP3A4.** A. 2D interaction diagram of CMP with HEM442, Ile301, Ser119, Leu482, Leu211, Ala305, Phe304 being closely interacting residues. B. 2D interaction diagram of Cyclosarin with HEM442, Ile301, Ser119, Leu482, Leu211, Ala305, Phe 304 being closely interacting residues. C. 2D interaction diagram of PiMP with HEM442, Ile301, Ala305, Phe 304, Ile369 and Ser119 being closely interacting residues. D. 2D interaction diagram of Soman with HEM442, Ile301, Ala305, Phe 304, Ile369 and Ser119 as interacting partners. E. 2D interaction diagram with HEM442, Thr309, Ser119, Phe304, Ile301, Ala305, Arg105 as major interacting residues with EMP. F. 2D interaction diagram of VX with HEM442, Thr309, Ser119, Phe304, Ile301, Ala305, Arg105 are major interacting residues. G. 2D interaction diagram of NEDPA HEM442, Thr309, Ser119, Phe304, Ile301, Ile369, Ala305 as major interacting residues. H. 2D interaction diagram of Tabun with HEM442, Thr309, Ser119, Phe304, Ile301, Ile369, Ala305 are major interacting residues. HEM – heme group in CYP3A4, Ser – Serine, Leu – Leucine, Ala – Alanine, Phe – Phenylalanine, Ile – Isoleucine, Thr – threonine, Arg – Arginine. All the interactions between CYP3A4 and OPs are given in interaction legend.

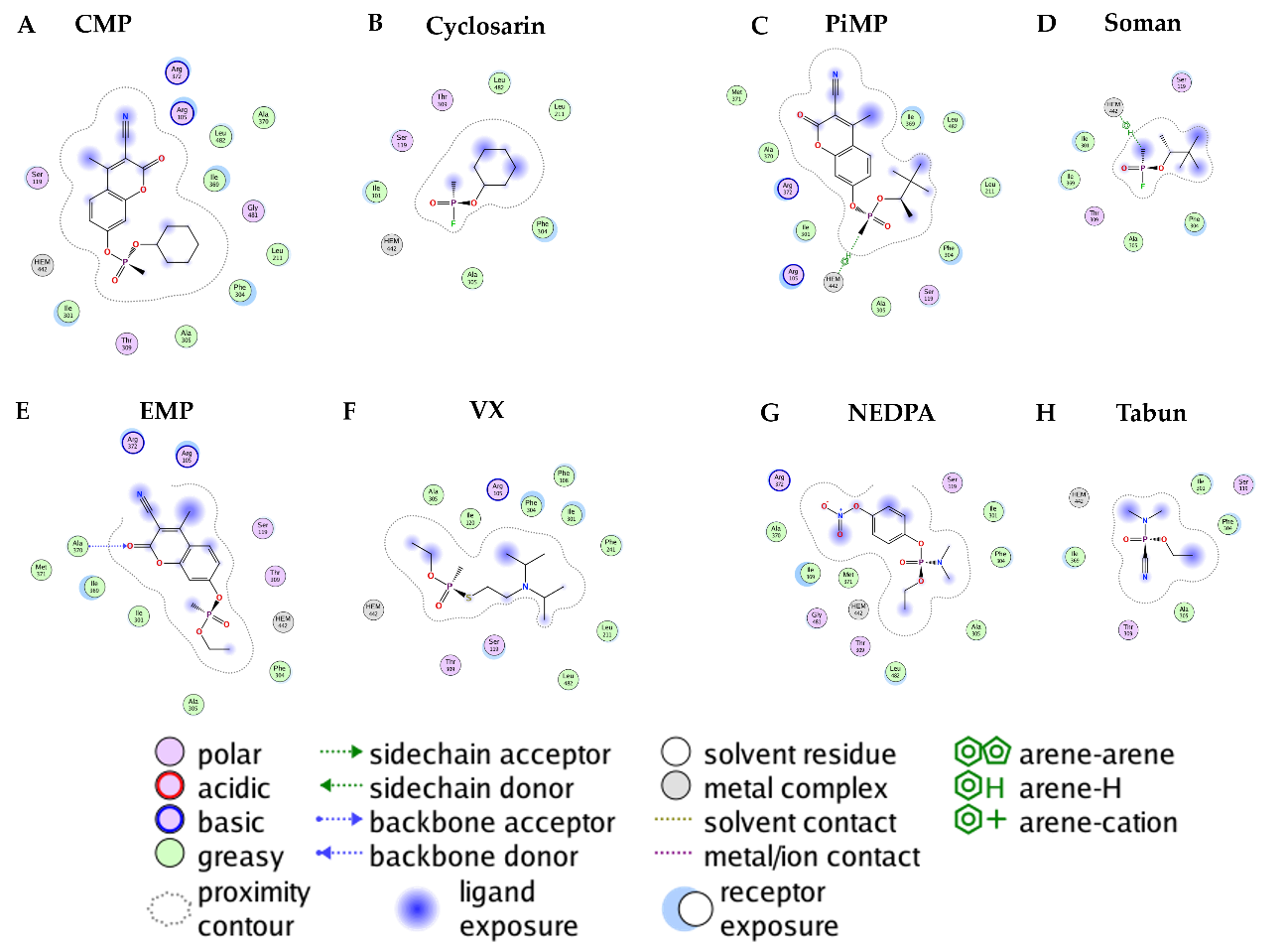

**Figure S6. 3D docking poses (shortest distance) for R and S conformations of EMP/VX and NEDPA/tabun with the 3NXU PDB structure of CYP3A4.** A. 3D docking poses for EMP (R) and VX (R). B. 3D docking poses for EMP (S) and VX (S). C. 3D docking poses for NEDPA (R) and Tabun (R). D. 3D docking poses for NEDPA (S) and Tabun (S). EMP is orange and Vx is cyan in color in panel A-B. NEDPA is orange and tabun is cyan in panel C-D.

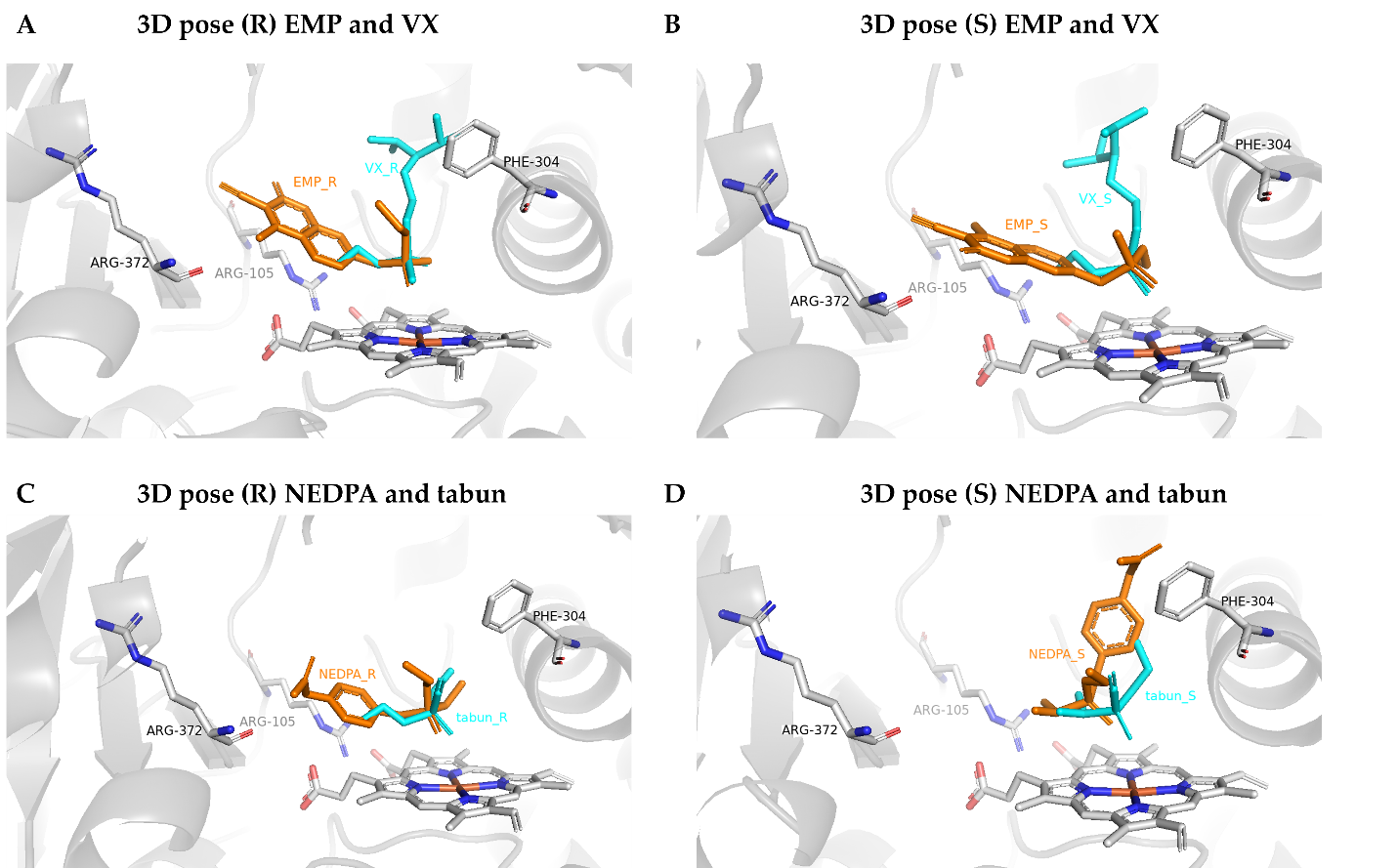

**Supplementary Tables**

**Supplementary Table 1: Table of molecular properties of the OPs.**

| **OP** | **Abbreviation** | **MW** | **LD_50_ (µM)** | **HBA** | **HBD** | **LogP** | **PSA** | **Rotatable bonds** |
| --- | --- | --- | --- | --- | --- | --- | --- | --- |
| Acephate | ACA | 183.17 | 163.88^38, 39^ | 2 | 1 | -0.001 | 55.4 | 4 |
| Azinphos-methyl | APM | 317.32 | 0.90^38, 39^ | 3 | 0 | 2.720 | 63.49 | 5 |
| Bensulide | BNS | 397.51 | 21.58^38, 39^ | 2 | 1 | 0.000 | 64.63 | 10 |
| Chlorfenvinphos | CFVP | 359.57 | 1.06^38, 39^ | 1 | 0 | 3.458 | 44.76 | 7 |
| Chlorphoxim | CPH | 332.74 | 238.66^38, 39^ | 2 | 0 | 4.688 | 63.84 | 8 |
| Chlorpyrifos | CPY | 350.59 | 5.98^38, 39^ | 1 | 0 | 4.967 | 40.05 | 6 |
| Chlorpyrifos Oxon | CPO | 334.50 | 6.27^38, 39^ | 2 | 0 | 4.233 | 57.12 | 6 |
| Cyclosarin surrogate | CMP | 361.33 | 0.32^38, 39^ | 3 | 0 | 3.016 | 85.62 | 5 |
| Crotoxyphos | CTP | 314.27 | -0.2^38, 39^ | 2 | 0 | 2.389 | 71.06 | 8 |
| Crufomate | CFA | 291.71 | 103.45^38, 39^ | 1 | 1 | 3.532 | 47.56 | 5 |
| Cyanofenphos | CFP | 303.30 | 4.50^38, 39^ | 1 | 0 | 4.452 | 42.25 | 6 |
| Diazinon | DIZ | 304.35 | 3.32^38, 39^ | 2 | 0 | 5.044 | 52.41 | 7 |
| Dichlorvos | DCV | 220.98 | 11.50^38, 39^ | 1 | 0 | 1.844 | 44.76 | 4 |
| Diethyl paraoxon | DEP | 275.19 | 0.35^38, 39^ | 2 | 0 | 1.697 | 96.57 | 7 |
| Diisopropyl fluorophosphate | DFP | 184.15 | 1.04^38, 39^ | 1 | 0 | 2.217 | 35.53 | 4 |
| Dimefox | DMX | 154.13 | 0.21^38, 39^ | 1 | 0 | 0.565 | 23.55 | 2 |
| Dimethoate | DMA | 229.26 | 33.95^38, 39^ | 1 | 1 | 0.445 | 47.56 | 6 |
| Dimethyl paraoxon | DMP | 247.14 | 0.26^38, 39^ | 2 | 0 | 0.975 | 96.57 | 5 |
| VX simulant | EMP | 307.24 | 0.38^33, 37^ | 3 | 0 | 1.773 | 85.62 | 5 |
| Ethoprophos | EPP | 242.34 | 5.24^38, 39^ | 1 | 0 | 3.206 | 26.3 | 8 |
| Fenamiphos | FMP | 303.36 | 0.63^38, 39^ | 1 | 1 | 3.401 | 47.56 | 7 |
| Fenthion | FNN | 278.33 | 3.64^38, 39^ | 0 | 0 | 4.090 | 27.69 | 5 |
| Formothion | FON | 257.30 | 30.86^38, 39^ | 2 | 0 | 0.452 | 55.84 | 7 |
| Iodofenphos | IFP | 413.00 | 179.21^38, 39^ | 0 | 0 | 5.160 | 27.69 | 4 |
| Iprobenfos | IBF | 288.34 | 74.91^38, 39^ | 1 | 0 | 4.186 | 35.53 | 7 |
| Isofenphos | IPP | 345.39 | 2.58^38, 39^ | 1 | 1 | 4.053 | 56.79 | 9 |
| Isoxathion | IXT | 313.31 | 11.36^38, 39^ | 2 | 0 | 4.307 | 49.28 | 7 |
| Leptophos | LPS | 412.06 | 3.32^38, 39^ | 0 | 0 | 5.902 | 18.46 | 4 |
| Malathion | MTN | 330.36 | 170.96^38, 39^ | 2 | 0 | 1.840 | 71.06 | 11 |
| Mephosfolan | MPF | 269.30 | 31.85^38, 39^ | 1 | 0 | 2.956 | 47.89 | 5 |
| Methamidophos | MMP | 141.13 | 6.75^38, 39^ | 1 | 1 | -0.660 | 52.32 | 2 |
| Methidathion | MDT | 302.33 | 2.63^38, 39^ | 3 | 0 | 3.205 | 60.36 | 6 |
| Mevinphos | MVP | 224.15 | 0.50^38, 39^ | 2 | 0 | 0.200 | 71.06 | 6 |
| Monocrotophos | MCP | 223.20 | 1.99^38, 39^ | 2 | 1 | 999.468 | 73.86 | 6 |
| Tabun surrogate | NEDPA | 274.21 | 27.52^32, 36^ | 2 | 0 | 1.699 | 90.58 | 6 |
| Phosalone | PHO | 367.81 | 10.36^38, 39^ | 1 | 0 | 4.427 | 48 | 7 |
| Phosfolan | PFN | 255.30 | 1.11^38, 39^ | 1 | 0 | 2.543 | 47.89 | 5 |
| Phosmet | PHT | 317.32 | 11.31^38, 39^ | 2 | 0 | 2.805 | 55.84 | 5 |
| Phosphamidon | PPM | 299.69 | 0.74^38, 39^ | 2 | 0 | 0.383 | 65.07 | 8 |
| Soman surrogate | PiMP | 363.50 | 0.97^32, 37^ | 3 | 0 | 3.563 | 85.62 | 6 |
| Pirimiphos ethyl | PPE | 333.39 | 13.34^38, 39^ | 3 | 0 | 4.500 | 55.65 | 9 |
| Pyrazophos | PZP | 373.36 | 12.85^38, 39^ | 4 | 0 | 3.456 | 81.95 | 9 |
| Pyridaphenthion | PPT | 340.33 | 71.78^38, 39^ | 2 | 0 | 1.836 | 60.36 | 7 |
| Quinalphos | QNP | 298.30 | 7.56^38, 39^ | 2 | 0 | 3.531 | 52.41 | 6 |
| Schradan | SCN | 286.25 | 0.56^38, 39^ | 2 | 0 | 0.716 | 56.33 | 6 |
| Temephos | TEM | 466.50 | 68.09^38, 39^ | 0 | 0 | 5.667 | 55.38 | 10 |
| Tetrachlorvinphos | TCVP | 365.95 | 347.20^38, 39^ | 1 | 0 | 4.897 | 44.76 | 5 |
| Triazophos | TAP | 313.31 | 6.69^38, 39^ | 3 | 0 | 3.781 | 55.65 | 7 |
| Tribufos | TBS | 314.50 | 3.12^38, 39^ | 1 | 0 | 5.207 | 17.07 | 12 |
| Tricholphon | TCP | 257.44 | 26.16^38, 39^ | 2 | 1 | 0.473 | 55.76 | 4 |

**Supplementary Table 2: Table of docking properties of the OPs in CYP3A4.**

| **OP** | **BE** | **top_score_dist** | **(BE_eff_)** | **(SD)** | **BE(SD)** | **BE_eff_(SD)** |
| --- | --- | --- | --- | --- | --- | --- |
| ACA | -4.25 | 8.152 | -0.024 | 2.961 | -4.05 | -0.023 |
| APM | -6.8 | 5.140 | -0.022 | 3.398 | -6.5 | -0.021 |
| BEN | -7.1 | 6.367 | -0.019 | 3.240 | -6.8 | -0.018 |
| CFP | -7.35 | 6.351 | -0.026 | 5.134 | -6.9 | -0.024 |
| CFVP | -6.75 | 5.675 | -0.019 | 4.010 | -6.25 | -0.018 |
| CMP | -8.15 | 5.460 | -0.023 | 4.330 | -7.65 | -0.023 |
| CPH | -6.7 | 5.959 | -0.021 | 3.712 | -6.7 | -0.021 |
| CPO | -6.2 | 10.370 | -0.019 | 3.365 | -6.2 | -0.019 |
| CPY | -6 | 6.207 | -0.018 | 3.441 | -5.8 | -0.017 |
| CFA | -6.8 | 3.079 | -0.025 | 3.079 | -6.8 | -0.025 |
| CTP | -7.3 | 6.690 | -0.025 | 3.620 | -6.9 | -0.023 |
| DCV | -4.7 | 6.594 | -0.022 | 2.678 | -4.5 | -0.021 |
| DFP | -5.1 | 4.430 | -0.030 | 3.301 | -4.7 | -0.028 |
| DIZ | -6.5 | 7.893 | -0.023 | 3.567 | -6.4 | -0.023 |
| DMX | -4 | 5.082 | -0.028 | 3.027 | -3.8 | -0.027 |
| DMA | -4.3 | 6.298 | -0.020 | 3.385 | -4.2 | -0.019 |
| EMP | -6.4 | 10.168 | -0.022 | 3.732 | -6.25 | -0.022 |
| EPP | -4.9 | 7.387 | -0.022 | 6.729 | -4.7 | -0.021 |
| FNN | -5.8 | 7.023 | -0.022 | 3.537 | -5.5 | -0.021 |
| FMP | -6.45 | 11.067 | -0.023 | 3.600 | -6.2 | -0.022 |
| IBF | -6.7 | 8.709 | -0.025 | 4.081 | -6.6 | -0.025 |
| IFP | -5.6 | 3.486 | -0.014 | 3.486 | -5.6 | -0.014 |
| IPP | -6.6 | 10.522 | -0.021 | 3.802 | -6.4 | -0.020 |
| IXT | -7.4 | 3.693 | -0.025 | 3.693 | -7.4 | -0.025 |
| LPS | -8 | 9.368 | -0.020 | 4.706 | -7.2 | -0.018 |
| MTN | -5.2 | 9.331 | -0.017 | 6.207 | -5 | -0.016 |
| MCP | -4.9 | 4.077 | -0.023 | 3.230 | -4.7 | -0.022 |
| MDT | -5.2 | 7.316 | -0.018 | 3.158 | -5.1 | -0.018 |
| MMP | -3.3 | 2.713 | -0.025 | 2.604 | -3.2 | -0.024 |
| MPF | -5.3 | 9.354 | -0.021 | 4.079 | -5.1 | -0.020 |
| MVP | -4.85 | 4.723 | -0.023 | 2.692 | -4.65 | -0.023 |
| NEDPA | -6.05 | 5.604 | -0.022 | 3.504 | -5.65 | -0.020 |
| PE | -6.2 | 11.098 | -0.024 | 2.740 | -6.1 | -0.023 |
| PHO | -6.5 | 6.298 | -0.018 | 4.421 | -6.2 | -0.018 |
| PHT | -7.3 | 5.597 | -0.024 | 3.627 | -7.2 | -0.024 |
| PiMP | -7.3 | 4.523 | -0.021 | 3.330 | -5.7 | -0.020 |
| PM | -5.6 | 4.490 | -0.024 | 2.737 | -5.6 | -0.024 |
| PPE | -6.2 | 8.339 | -0.020 | 6.177 | -6 | -0.019 |
| PPM | -5.6 | 5.238 | -0.020 | 4.343 | -5.5 | -0.020 |
| PPT | -7.5 | 3.654 | -0.023 | 3.542 | -7.1 | -0.022 |
| PZP | -6.5 | 11.507 | -0.018 | 3.375 | -5.9 | -0.017 |
| QPS | -6.9 | 4.899 | -0.024 | 3.470 | -6.5 | -0.023 |
| SCN | -4.8 | 8.740 | -0.018 | 4.183 | -4.5 | -0.017 |
| TAP | -7.2 | 3.484 | -0.024 | 3.484 | -7.2 | -0.024 |
| TCP | -4.5 | 6.035 | -0.018 | 4.044 | -4.3 | -0.017 |
| TCVP | -6.55 | 6.535 | -0.018 | 5.135 | -6.3 | -0.017 |
| TBS | -5.9 | 8.668 | -0.021 | 4.570 | -5.4 | -0.019 |

BE – Binding Energy, BEeff – Binding efficiency, D – distance between heme center in CYP3A4 and phosphorous atom in OP for top pose, SD – shortest distance between heme center in CYP3A4 and phosphorous atom in OP

**Supplementary Table 3. Correlation between the time-dependent difference in inhibition of recombinant CYP3A4 at 1 µM (10 and 30 minutes) with the molecular parameters.**

| **Property** | **Pearson R** | **P (two-tailed)** | **Slope** | **Y-intercept** |
| --- | --- | --- | --- | --- |
| *MW | -0.359 | 0.013 | -0.565 | 293.300 |
| LD50 (µM) | -0.017 | 0.910 | -0.031 | 34.770 |
| HBA | -0.272 | 0.065 | -0.006 | 1.639 |
| HBD | 0.255 | 0.083 | 0.003 | 0.218 |
| LogP | -0.170 | 0.255 | -0.007 | 2.708 |
| PSA | -0.196 | 0.187 | -0.097 | 54.510 |
| Rotatable bonds | -0.103 | 0.492 | -0.005 | 6.200 |
| *VWR (Å) | -0.310 | 0.034 | -0.430 | 254.800 |
| *BE | 0.354 | 0.015 | 0.010 | -5.965 |
| D | 0.119 | 0.426 | 0.007 | 6.741 |
| BE_eff_ | -0.134 | 0.368 | -1.05E-05 | -0.022 |
| SD | -0.116 | 0.437 | -0.003 | 3.765 |
| *BE(SD) | 0.305 | 0.037 | 0.008 | -5.722 |
| BE_eff_(SD) | -0.130 | 0.385 | -9.85E-06 | -0.021 |

Parameters with * are significantly correlated with experimental result

**Supplementary Table 4. Correlation between the time-dependent difference in inhibition of recombinant CYP3A4 at 10 µM (10 and 30 minutes) with molecular parameters.**

| **Property** | **Pearson R** | **P (two-tailed)** | **Slope** | **Y-intercept** |
| --- | --- | --- | --- | --- |
| *MW | 0.321 | 0.028 | 0.474 | 295.000 |
| LD50 (µM) | 0.220 | 0.138 | 0.370 | 31.820 |
| HBA | -0.040 | 0.788 | -0.001 | 1.710 |
| HBD | 0.091 | 0.541 | 0.001 | 0.184 |
| LogP | 0.248 | 0.093 | 0.010 | 2.696 |
| PSA | -0.090 | 0.547 | -0.042 | 55.880 |
| Rotatable bonds | 0.150 | 0.315 | 0.007 | 6.192 |
| VWR (Å) | 0.240 | 0.105 | 0.311 | 256.500 |
| BE | -0.078 | 0.603 | -0.002 | -6.050 |
| D | -0.055 | 0.714 | -0.003 | 6.695 |
| *BE_eff_ | 0.349 | 0.016 | 2.55E-05 | -0.022 |
| SD | 0.106 | 0.478 | 0.002 | 3.773 |
| BE(SD) | -0.081 | 0.590 | -0.002 | -5.789 |
| BE_eff_(SD) | 0.334 | 0.022 | 2.38E-05 | -0.021 |

Parameters with * are significantly correlated with experimental result.

**Supplementary Table 5. Correlation between the inhibition of recombinant CYP3A4 at 1 µM at 10 minutes with the molecular parameters.**

| **Property** | **Pearson R** | **P (two-tailed)** | **Slope** | **Y-intercept** |
| --- | --- | --- | --- | --- |
| MW | -0.139 | 0.351 | -0.191 | 301.000 |
| LD50 (µM) | -0.070 | 0.638 | -0.110 | 36.130 |
| HBA | -0.074 | 0.619 | -0.001 | 1.716 |
| HBD | 0.097 | 0.515 | 0.001 | 0.184 |
| LogP | -0.123 | 0.409 | -0.005 | 2.830 |
| PSA | -0.034 | 0.820 | -0.015 | 55.650 |
| Rotatable bonds | -0.006 | 0.966 | -2.9E-03 | 6.258 |
| VWR (Å) | -0.096 | 0.520 | -0.116 | 260.400 |
| BE | 0.141 | 0.345 | 0.003 | -6.101 |
| D | 0.090 | 0.547 | 0.005 | 6.624 |
| BE_eff_ | -0.034 | 0.822 | -2.29E-06 | -0.022 |
| SD | -0.159 | 0.287 | -0.003 | 3.823 |
| BE(SD) | 0.104 | 0.488 | 0.002 | -5.829 |
| BE_eff_(SD) | -0.024 | 0.873 | -1.59E-06 | -0.021 |

Parameters with * are significantly correlated with experimental result; Pearson Correlation.

**Supplementary Table 6. Correlation between the inhibition of recombinant CYP3A4 at 1 µM at 30 minutes with the molecular parameters.**

| **Property** | **Pearson R** | **P (two-tailed)** | **Slope** | **Y-intercept** |
| --- | --- | --- | --- | --- |
| MW | 0.257 | 0.081 | 0.523 | 288.800 |
| LD50 (µM) | -0.082 | 0.583 | -0.191 | 38.880 |
| HBA | 0.241 | 0.103 | 0.007 | 1.565 |
| HBD | -0.185 | 0.212 | -0.002 | 0.239 |
| LogP | 0.036 | 0.808 | 0.002 | 2.745 |
| PSA | 0.203 | 0.172 | 0.129 | 52.950 |
| Rotatable bonds | 0.123 | 0.410 | 0.008 | 6.094 |
| VWR (Å) | 0.258 | 0.080 | 0.462 | 250.100 |
| BE | -0.249 | 0.092 | -0.009 | -5.889 |
| D | -0.020 | 0.893 | -0.002 | 6.698 |
| BE_eff_ | 0.123 | 0.408 | 1.24E-05 | -0.022 |
| SD | -0.085 | 0.570 | -0.002 | 3.842 |
| BE(SD) | -0.241 | 0.103 | -0.008 | -5.642 |
| BE_eff_(SD) | 0.132 | 0.377 | 1.29E-05 | -0.021 |

Parameters with * are significantly correlated with experimental result; Pearson Correlation.

**Supplementary Table 7. Correlation between inhibition of recombinant CYP3A4 at 10 µM 30 minutes with molecular parameters**

| **Property** | **Pearson R** | **P (two-tailed)** | **Slope** | **Y-intercept** |
| --- | --- | --- | --- | --- |
| *MW | 0.690 | <0.0001 | 0.982 | 244.200 |
| LD50 (µM) | 0.267 | 0.069 | 0.302 | 18.180 |
| HBA | 0.160 | 0.283 | 0.003 | 1.508 |
| HBD | -0.227 | 0.125 | -0.003 | 0.335 |
| *LogP | 0.569 | <0.0001 | 0.024 | 1.452 |
| PSA | -0.054 | 0.721 | -0.057 | 58.710 |
| *Rotatable bonds | 0.333 | 0.022 | 0.017 | 5.280 |
| *VWR (Å) | 0.597 | <0.0001 | 0.792 | 215.000 |
| *BE | -0.484 | 0.001 | -0.013 | -5.315 |
| D | -0.042 | 0.778 | -0.003 | 6.830 |
| *BE_eff_ | 0.292 | 0.047 | 2.56E-05 | -0.023 |
| SD | 0.192 | 0.196 | 0.004 | 3.557 |
| *BE(SD) | -0.443 | 0.002 | -0.012 | -5.122 |
| *BE_eff_(SD) | 0.312 | 0.033 | 2.52E-05 | -0.022 |

Parameters with * are significantly correlated with experimental result; Spearman Correlation.

**Supplementary Table 8. Correlation between inhibition of recombinant CYP3A4 at 10 µM 10 minutes with molecular parameters.**

| **Property** | **Pearson R** | **P (two-tailed)** | **Slope** | **Y-intercept** |
| --- | --- | --- | --- | --- |
| *MW | 0.423 | 0.003 | 0.474 | 276.800 |
| LD50 (µM) | -0.044 | 0.771 | -0.076 | 38.650 |
| HBA | 0.184 | 0.217 | 0.004 | 1.506 |
| *HBD | -0.359 | 0.013 | -0.003 | 0.347 |
| *LogP | 0.349 | 0.016 | 0.013 | 2.182 |
| PSA | 0.018 | 0.903 | -0.014 | 56.160 |
| Rotatable bonds | 0.207 | 0.162 | 0.010 | 5.802 |
| *VWR (Å) | 0.402 | 0.005 | 0.453 | 237.900 |
| *BE | -0.453 | 0.001 | -0.011 | -5.555 |
| D | 0.038 | 0.798 | 2.2E-03 | 6.658 |
| BE_eff_ | 0.079 | 0.596 | 6.47E-07 | -0.022 |
| SD | 0.046 | 0.761 | 0.002 | 3.707 |
| *BE(SD) | -0.413 | 0.004 | -0.010 | -5.347 |
| BE_eff_(SD) | 0.095 | 0.527 | 6.76E-07 | -0.021 |

Parameters with * are significantly correlated with experimental result; Spearman Correlation.

**Supplementary Table 9. Correlation between LC-MS/MS 2 hours metabolism by recombinant CYP3A4 with molecular parameters.**

| **Property** | **Pearson R** | **P (two-tailed)** | **Slope** | **Y-intercept** |
| --- | --- | --- | --- | --- |
| *MW | -0.455 | 0.033 | -0.739 | 332.900 |
| LD50 (µM) | 0.283 | 0.202 | 0.624 | 18.300 |
| HBA | -0.086 | 0.704 | -0.002 | 1.703 |
| HBD | 0.029 | 0.899 | 2.3E-03 | 0.170 |
| *LogP | -0.446 | 0.038 | -0.020 | 3.594 |
| PSA | 0.135 | 0.548 | 0.034 | 52.100 |
| *Rotatable bonds | -0.476 | 0.025 | -0.023 | 7.608 |
| *VWR (Å) | -0.565 | 0.006 | -0.841 | 294.700 |
| *BE | 0.573 | 0.005 | 0.016 | -6.580 |
| D | -0.038 | 0.867 | -0.002 | 6.954 |
| BE_eff_ | 0.014 | 0.951 | 1.11E-06 | -0.021 |
| SD | -0.126 | 0.576 | -0.005 | 4.409 |
| *BE(SD) | 0.577 | 0.005 | 0.016 | -6.342 |
| BE_eff_(SD) | 0.041 | 0.857 | 3.59E-06 | -0.020 |

Parameters with * are significantly correlated with experimental result; Pearson Correlation.

**Supplementary Table 10. Correlation between LC-MS/MS 1 hour metabolism by recombinant CYP3A4 with molecular parameters.**

| **Property** | **Pearson R** | **P (two-tailed)** | **Slope** | **Y-intercept** |
| --- | --- | --- | --- | --- |
| MW | -0.359 | 0.072 | -0.534 | 331.500 |
| LD50 (µM) | 0.203 | 0.320 | 0.399 | 19.410 |
| HBA | -0.155 | 0.451 | -3.0E-03 | 1.952 |
| HBD | -0.032 | 0.876 | 0.000 | 0.168 |
| *LogP | -0.433 | 0.027 | -0.017 | 3.553 |
| PSA | 0.190 | 0.354 | 0.072 | 55.270 |
| Rotatable bonds | -0.365 | 0.067 | -0.016 | 7.244 |
| *VWR (Å) | -0.448 | 0.022 | -0.630 | 295.500 |
| *BE | 0.461 | 0.018 | 0.013 | -6.684 |
| D | 0.053 | 0.796 | 0.003 | 6.654 |
| BE_eff_ | 0.079 | 0.701 | 5.64E-06 | -0.021 |
| SD | -0.016 | 0.938 | -0.001 | 4.153 |
| BE(SD) | 0.473 | 0.015 | 0.013 | -6.367 |
| BE_eff_(SD) | 0.115 | 0.574 | 9.11E-06 | -0.021 |

Parameters with * are significantly correlated with experimental result; Pearson Correlation.

**Supplementary Table 11 Correlation P-values between molecular parameters. Pearson correlation was used.**

|  | **MW** | **LD50** | **HBA** | **HBD** | **LogP** | **PSA** | **RRB** | **VWR** | **BE** | **D** | **BE_eff_** | **SD** | **BE**  **(SD)** | **BE_eff_**  **(SD)** |
| --- | --- | --- | --- | --- | --- | --- | --- | --- | --- | --- | --- | --- | --- | --- |
| **MW** |  | NS | NS | NS | **** | NS | *** | **** | **** | NS | **** | * | **** | **** |
| **LD50** | NS |  | NS | NS | NS | NS | NS | NS | NS | NS | NS | NS | NS | NS |
| **HBA** | NS | NS |  | NS | NS | **** | NS | NS | NS | NS | NS | NS | NS | NS |
| **HBD** | NS | NS | NS |  | ** | NS | NS | NS | * | NS | NS | NS | * | NS |
| **LogP** | **** | NS | NS | ** |  | ** | NS | **** | **** | NS | NS | * | **** | NS |
| **PSA** | NS | NS | **** | NS | ** |  | NS | NS | NS | NS | NS | NS | NS | NS |
| **RRB** | *** | NS | NS | NS | NS | NS |  | **** | * | ** | * | ** | * | * |
| **VWR** | **** | NS | NS | NS | **** | NS | **** |  | **** | NS | ** | ** | **** | ** |
| **BE** | **** | NS | NS | * | **** | NS | * | **** |  | NS | NS | NS | **** | NS |
| **D** | NS | NS | NS | NS | NS | NS | ** | NS | NS |  | NS | NS | NS | * |
| **BE_eff_** | **** | NS | NS | NS | NS | NS | * | ** | NS | NS |  | * | NS | **** |
| **SD** | * | NS | NS | NS | * | NS | ** | ** | NS | NS | * |  | NS | * |
| **BE**  **(SD)** | **** | NS | NS | * | **** | NS | * | **** | **** | NS | NS | NS |  | NS |
| **BE_eff_**  **(SD)** | **** | NS | NS | NS | NS | NS | * | ** | NS | * | **** | * | NS |  |
